# Transcriptional Dosage of Oncogenic KRAS Drives Lung Adenocarcinoma Cell States, Progression and Metastasis

**DOI:** 10.1101/2024.12.29.630643

**Authors:** Michela Serresi, Ali Osman Cetin, Yuliia Dramaretska, Sonia Kertalli, Matthias J. Schmitt, Heike Naumann, Maria Zschummel, Marie Liesse-Labat, Lucas F. Maciel, Jeroen Declercq, Jean-Christophe Marine, Gaetano Gargiulo

## Abstract

Cancer cells display distinct, recurrent phenotypic cell states. Metastatic spreading correlates with tumor cell state evolution. However, the molecular mechanisms underlying metastasis remain elusive. Here, we demonstrate that the quantitative dosage of oncogenic KRAS drives lung adenocarcinoma progression and metastasis via the integration of external signaling and pioneer transcription factor dynamics into qualitative cell states. Combining mouse models, in vivo CRISPR activation screens, and fate mapping, we show that even mild transcriptional amplification of KRAS significantly fuels tumor progression and metastasis. Chromatin profiling and transcriptomics reveal that high and low KRAS dosages supersede and integrate inflammatory and TGFβ signaling to dictate mouse cancer cell states. Patient data show that KRAS dosages correlate with distinct survival outcomes, transcription factor activity, and cell states. Direct KRAS inhibition in xenografts limits the KRAS-high “proliferative” cell state but spares a minimal residual state mimicking the KRAS-low “ciliated-like” state. Thus, oncogenic KRAS dosage fuels tumor heterogeneity at the cell state level and drives a bimodal tumor evolution during metastasis, with implications for prognosis and treatment.

## Introduction

Metastasis, the spread of cancer cells from a primary to distant organs, significantly increases mortality risk among cancer patients ^1–3^. Genetically, cancer drivers in metastatic tumors often mirror those of their primary counterpart ^4,5^, indicating that additional genetic alterations may not be essential for tumor progression ^6^. Metastasis involves limited copy number aberrations, which may influence therapeutic responses but do not fully explain metastatic spread ^4^. The distribution of tumor microenvironments (TMEs) is largely consistent across most primary and metastatic cancers ^4^, which might support an hourglass evolution model by which non-genetic factors could critically influence cancer cell fates during progression and metastasis. In certain contexts, ‘driver cell states,’ may supersede genetic alterations in driving tumor initiation, progression, therapeutic response, and metastasis ^7^. How driver genetics and driver cell states are connected remains to be systematically investigated and is likely context-dependent.

RAS proteins, among the most intensively studied oncogenes in cancer biology, became recently ‘druggable’ ^8–11^. Directly targeting KRAS through mutation-specific and pan-active forms is expected to extend patient survival but will also increase metastatic recurrence. Given this, it is timely to expand focus on how RAS proteins influence cancer progression and metastasis. KRAS mutations are present in approximately 32% of non-small cell lung cancer (NSCLC) cases, the most prevalent cancer globally, particularly affecting the lung adenocarcinoma (LUAD) subtype ^12^.

*KRAS* fits well the definition of a prototypic driver gene in human cancer ^13^. *KRAS* mutations in LUAD and pancreatic adenocarcinoma (PAAD) often represent founding mutations, whereas in the context of Colorectal Adenocarcinoma (COAD), *KRAS* mutant typically act more as a “promoter” ^14^. In all the cases, mutations in *KRAS* are associated with poor prognosis, not least due to the lack of therapeutics until recently. Autochthonous mouse models for KRAS-driven cancers like LUAD and PAAD closely mimic human disease ^15,16^, offering predictive value for therapeutic responses ^17^. In the most prevalent KRAS-driven mouse model, a Kras^G12D^ allele is activated through genetic recombination, resulting in heterozygous oncogenic expression in the targeted tissues ^15,16^. In such models, metastatic events are infrequent, but the progression is driven by the collaboration between Kras^G12D^ and the deletion of tumor suppressor genes such as Trp53, Lkb1 and Eed, as well as non-genetic tumor evolution ^18–22^. In mouse LUAD, oncogenic Kras can transform multiple cell types ^23^ and displays a broad range of cell states ranging from those mirroring the candidate cell of origin towards evolving a mesenchymal-like state enriched at late stages and metastasis ^18^. While KRAS-associated metabolism and microenvironment are key players in shaping progression^24,25^, how cell state evolution occurs in these models remains unclear.

Intriguingly, in Kras-driven mouse PDAC, copy number aberrations increasing the genetic dosage of the hyperactive Kras oncogene were observed ^26^. Moreover, allelic imbalance between wild-type and oncogenic KRAS drives metabolic rewiring ^27^. This suggests that Kras mutations and dosage serve two distinct non-overlapping roles in tumor progression. However, the need to deduce transcriptional or protein dosage from copy numbers, combined with the early occurrence of such events, complicates the assessment of their impact on progression and metastasis. Transplantation-based models provide practical advantages over the rapid progression of aggressive LUAD and the impracticality of surgical resection in autochthonous models. These models offer unique opportunities for elaborate genetic manipulations and challenging *in vivo* genetic screens ^28,29^ and the relevance of findings generated in these models may be augmented by data analyses of large repositories of cellular and molecular cancer obtained in an intact mouse and human cancers.

In this study, we investigated the molecular mechanisms driving Kras-driven LUAD progression and metastasis, focusing on a model for mucinous LUAD—an advanced stage driven by intrinsic cancer cell state evolution^20,30^ and associated with resistance to KRAS inhibition^31^. To this end, we used the Kras^G12D^;Trp53^-/-^;Eed^-/-^ (KPE) genotype, which recapitulates these features in mice ^20^. KPE cells uniformly progress toward pre-metastatic stages and retain high-grade phenotypes after transplantation, making this setting ideal for studying metastasis via CRISPR activation and fate mapping screens. By integrating findings across KPE and other model systems (e.g., KP mouse models and human xenografts), and testing the generalizability of our findings in human primary and metastatic LUAD transcriptional repositories, we establish KRAS dosage as a robust driver of metastatic dynamics dictating global chromatin and transcriptional changes ultimately influencing cancer cell states and progression, with broad implications for metastasis and therapeutic responses.

## Results

### Fate mapping and Transcriptional amplification during lung adenocarcinoma progression and metastasis by cellular barcoding and CRISPR activation *in vivo*

We previously established mouse models for aggressive LUAD initiation, progression and metastasis ^20,28^. To study whether modulating endogenous gene expression levels of cancer genes in the lung microenvironment drives functional outcomes during murine LUAD progression and metastasis, we set up an *in vivo* screening platform using an orthotopic setting. The first component of such *in vivo* platform consists of primary cells derived from autochthonous lung tumor model capable of disease-relevant transplantation. We used KPE primary cells from mice with a KRAS^G12Dfl/+^;Trp53^fl/fl^;Eed^fl/fl^ background, derived from LUAD induced by adenoviral-CRE. Compared to the most commonly used KP genotype, this model homogeneously progresses towards a pre-metastatic stage *in vivo* via EMT and phenotypic switching towards mucinous differentiation ^20^. The latter characteristic is cell intrinsic and retained upon transplantation in secondary recipients ^20^. A second component of this platform is a CRISPRa system with a doxycycline-inducible dCas9 fused to a VPR trans-activator ^32^, and a library of gRNAs cloned in a CROP-seq vector ^33^. We chose gRNA targeting 88 genes, enriched in those functionally driving progression in a transplantation model for LUAD progression, such as Kras and NF-kB/PRC2 targets ^28^. A set of ∼10 /gene on-target gRNAs was complemented with an equal number of non-targeting gRNAs. Together, the combined system enables transcriptional amplification of target genes via locus-specific gRNAs or population-level cellular fate mapping via non-targeting gRNAs serving as neutral barcodes (**Fig. 1A)**. Detection of *in vivo* bioluminescence via firefly Luciferase expression enables time-resolved lung grafting prior to gRNA activation (**Fig. S1A)**.

**Figure 1.**
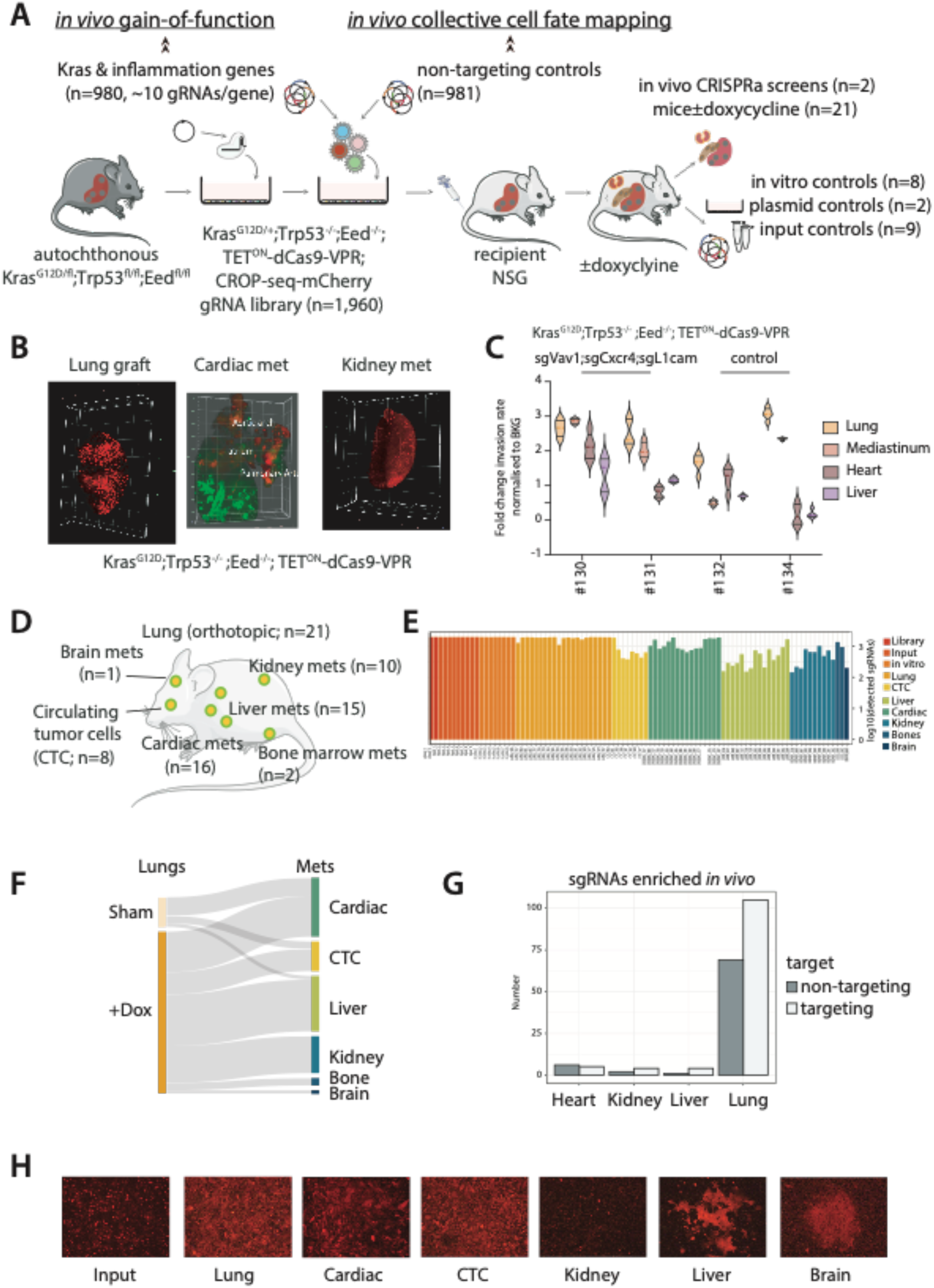
In vivo gain-of-function and fate mapping of Kras-driven lung cancer cells by CRISPRa-CROP-seq. A) Experimental outline: Kras^G12D^;Trp53 ^-/-^;Eed^-/-^;Tet_ON_-dCas9-VPR (KPE-VPR) were low-MOI- infected with a lentiviral library of gRNAs targeting Kras-associated inflammatory mediators (from Serresi et al., 2018) and non-targeting gRNAs serving as barcodes for neutral evolution. KPE-VPR; CROP-mCherry were orthotopically transplanted into recipient mouse lungs and gRNA expression was activated *in vivo* by doxycycline upon grafting was verified (see Fig.S1). Plasmid library, input and *in vitro* expanded cells served as control for stochastic gRNAs drift. B) Representative lightsheet microscopy of lung, heart and kidney from mice transplanted with the cells indicated background. C) Violin plot of bioluminescence (BLI) emission at the humane-end-point from *ex-vivo* isolated organs from mice injected with the indicated KPE cells. Regions of interest (ROIs) were guided by BLI signal and normalized over background-positioned ROIs. D) Graphical summary of the tissue samples collected in the screenings. E) Bar plot of sgRNA detection rate from all the samples in the screen. Color code denotes cells or tissue of origin. F) Sankey plots showing the tumor evolution in animals treated with doxycycline-inducing gRNA library activation, or sham (no gRNA activation). G) Bar plot showing the linear counts of non-targeting and targeting sgRNA value in primary lung tumors and in representative distant organs. H) Representative images of isolated primary passage 0 cancer cells from different organs. mCherry marks gRNAs containing cancer cells.

To test whether this system is well versed to study whether transcriptional amplification of candidates oncogenes impacts progression, we concomitantly delivered to the same cells three gRNA, each targeting endogenous *Vav1*, *Cxcr4* and *L1cam*. These were previously validated *in vivo* using standard viral-driven ORF amplification^28^. Compared to control cells, dox-treatment induced significant transcriptional and protein expression of the VPR-dCas9 in a time window of 20 days, that was maintained after dox-washout, and promoted up to 40-fold transcriptional amplification of *Vav1*, *Cxcr4* and *L1cam* (**Fig. S1B-C)**. KPE cells colonized both lung and several organs of recipient NSG mice as individual small cell clusters as gauged by ex vivo light sheet microscopy of CUBIC cleared tissues. Together with the increased tumor burden in presence of targeted transcriptional amplification, both support the robustness of the platform (**Fig. 1B-C)**.

To run simultaneous clonal analysis of the impact of transcriptional amplification of single genes and population-level fate mapping of KPE dynamics during progression and metastasis, we next synthesized a library of 1,981 gRNAs equally divided between targeting and 981 non-targeting gRNAs and planned a theoretical representation at the time of chemogenetic library activation of >1,000 gRNAs per tumor, assuming a single infection per cell ^29^. We performed two independent screens and used ex vivo bioluminescence to guide dissection of tissues carrying substantial primary or metastatic burdens (**Fig. 1A and S1**). Compared to plasmid and input controls, massively parallel library preparation and sequencing in passage zero cell culture from each tissue in each mouse confirmed the overall broad representation of input gRNA in orthotopic tumors (**Fig. 1D-G** and **S1D-E)** and revealed its progressive reduction in metastatic lesions (**Fig. 1E** and **S1E)**. Dox-activation selectively increased targeting gRNA within the context of a uniform segregation of targeting and non-targeting gRNAs across tissues (**Fig. S1F-G**). This indicates that transcriptional amplification of oncogenes is not required for metastasis in the KPE model but can increase the metastatic rate. Of note, tissue-derived tumor cells display phenotypic differences and variable EMT biomarkers (**Fig. 1H** and **S1G-I**), likely the result of distinct and heritable phenotypic adaptation in tissues.

Overall, our data support that coupling fate mapping and CRISPR activation *in vivo* enables the functional dissection of genes regulating progression and metastasis in a mouse model for aggressive NSCLC.

### Kras transcriptional amplification promotes oligo-/poly-clonal orthotopic and metastatic growth in a Kras^G12D^-driven lung adenocarcinoma

To analyse our *in vivo* screens in a statistically robust manner, we devised a strategy that shortlisted homogeneously diverse samples, which builds on key parameters such as the number of reads per guide and inter-sample correlation (**Fig. 2A**, **S2A-C** and methods). This strategy enabled us to compare and contrast input, orthotopic and metastatic samples that are quantitatively comparable. In turn, this approach uncovered that orthotopic samples represent both targeting and non-targeting gRNAs in Gaussian fashion as predicted by light sheet imaging (**Fig. 1B)** and a bimodal gRNA distribution defines oligoclonal and polyclonal metastases as the prevalent mode of dissemination in the KPE model (**Fig. 2B)**.

**Figure 2.**
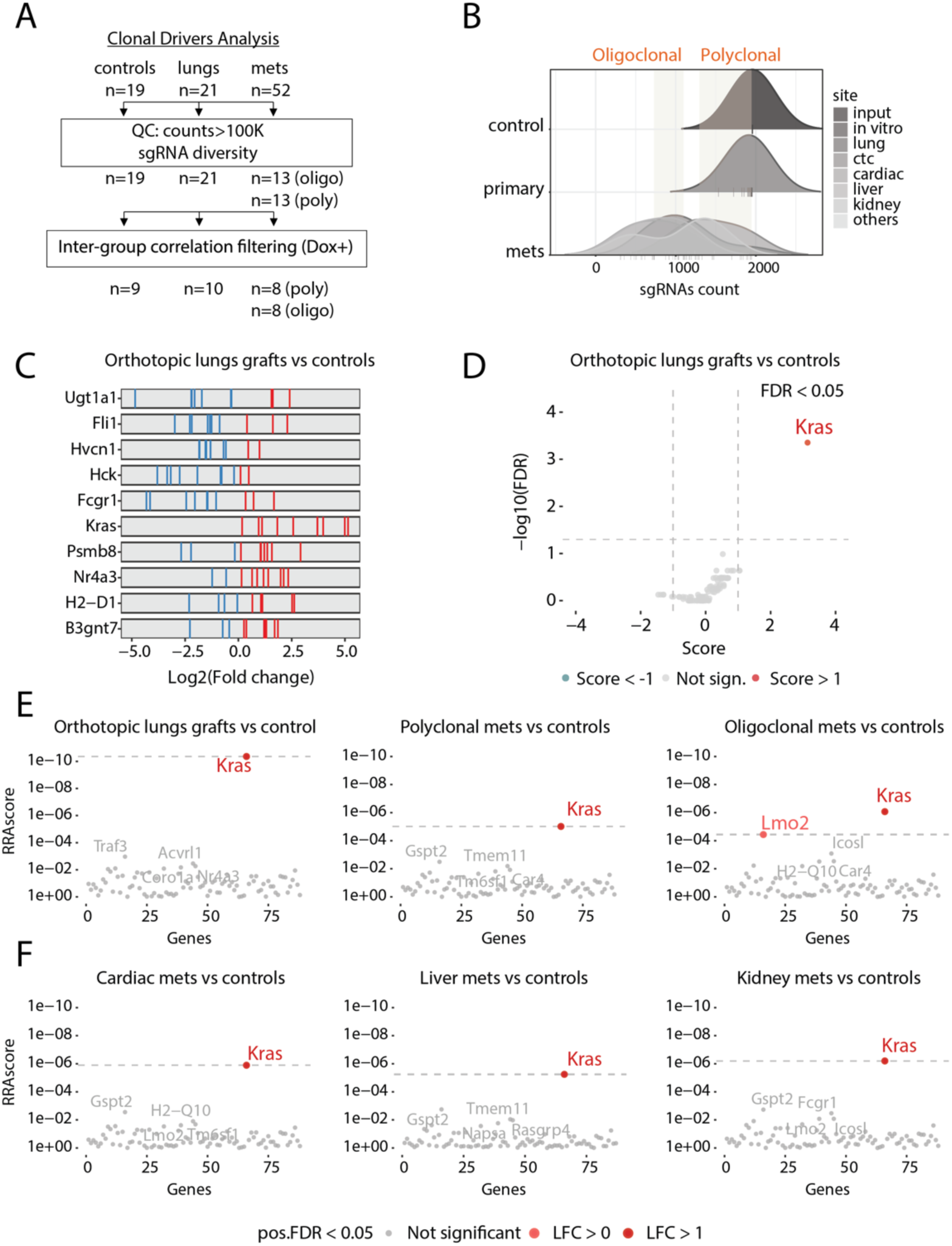
Kras transcriptional amplification promotes orthotopic and metastatic growth in a KrasG12D-driven lung cancer. **A)** Schematics of quality control (QC) pipeline. **B)** Density distribution plot of the total gRNAs retrieved in each sample. Oligoclonal and Polyclonal metastases were defined as 25-50 and >75 percentile of frequency. **C)** Visualization of enrichment/depletion scoring for gRNAs targeting a selected subset of genes compared to non-targeting controls; for each gene, red and blue denote gRNAs with fold-change >0 and <0, respectively. **D)** Volcano plot showing False Discovery Rate (FDR) significance for orthotopic lung grafts at the gene level. **E**-**F**) MAGeCK plot showing Robust Rank Aggregation (RRA) scores for all genes in the library in core screen samples divided into the indicated classes (see Fig. S2 and methods). FDR significance threshold denotes Kras as the key gene for orthotopic and metastatic growth.

Next, we applied two parallel approaches to nominate the driver(s) among the genes tested *in vivo* (methods). This approach showed that Kras was the sole gene within our pool to systematically reach statistical significance (**Fig. 2E-F** and **S2C-D**). In turn, this reinforces that a KPE genotype is fully penetrant for both tumor initiation, progression and metastasis in cell intrinsic manner (e.g. in a immunodeficient model for tumor progression). Of note, the cardiac vasculature appeared to host highly polyclonal KPE cells. Ranking RRA scores and focusing on individual tissues opened to the potential selection of additional hits, such as *IcosL* and *Napsa* in the Kidney and Liver setting (**Fig. 2F** and **S2D**), but *Kras* remains the top hit. Surprisingly, well-established oncogenes such as *Vav1*, *Cxcr4* and *L1cam* previously validated via overexpression of human open-reading frames using strong viral promoters did not score as hits despite their dosage was amplified via their endogenous locus ^28^.

To validate Kras as the single event promoting mouse LUAD, we set up independent validation experiments. First, we set up an *in vivo* competition assay, in which seven top ranking gRNAs targeting endogenous *Kras* transcriptional amplification were pooled and low-MOI transduced KPE;dCas9-VPR cells with mCherry fluorescence marker. These cells were mixed in an unfavorable ratio with KPE;dCas9-VPR cells infected with a large library of gRNAs targeting the human-genome (n=104,504), which are therefore off-target in the mouse (**Fig.3A-B)**. In all organs in which bioluminescence highlighted substantial tumor burden, transcriptional amplification of Kras drove competitive advantage in both short *in vitro* assay with a single Kras guide, as well as during tumor transplantation *in vivo*, as gauged by mCherry-labeled cells bearing seven distinct on-target guides outnumbering neutral guide-bearing BFP-labeled ones (**Fig. 3A** and **3C**).

**Figure 3.**
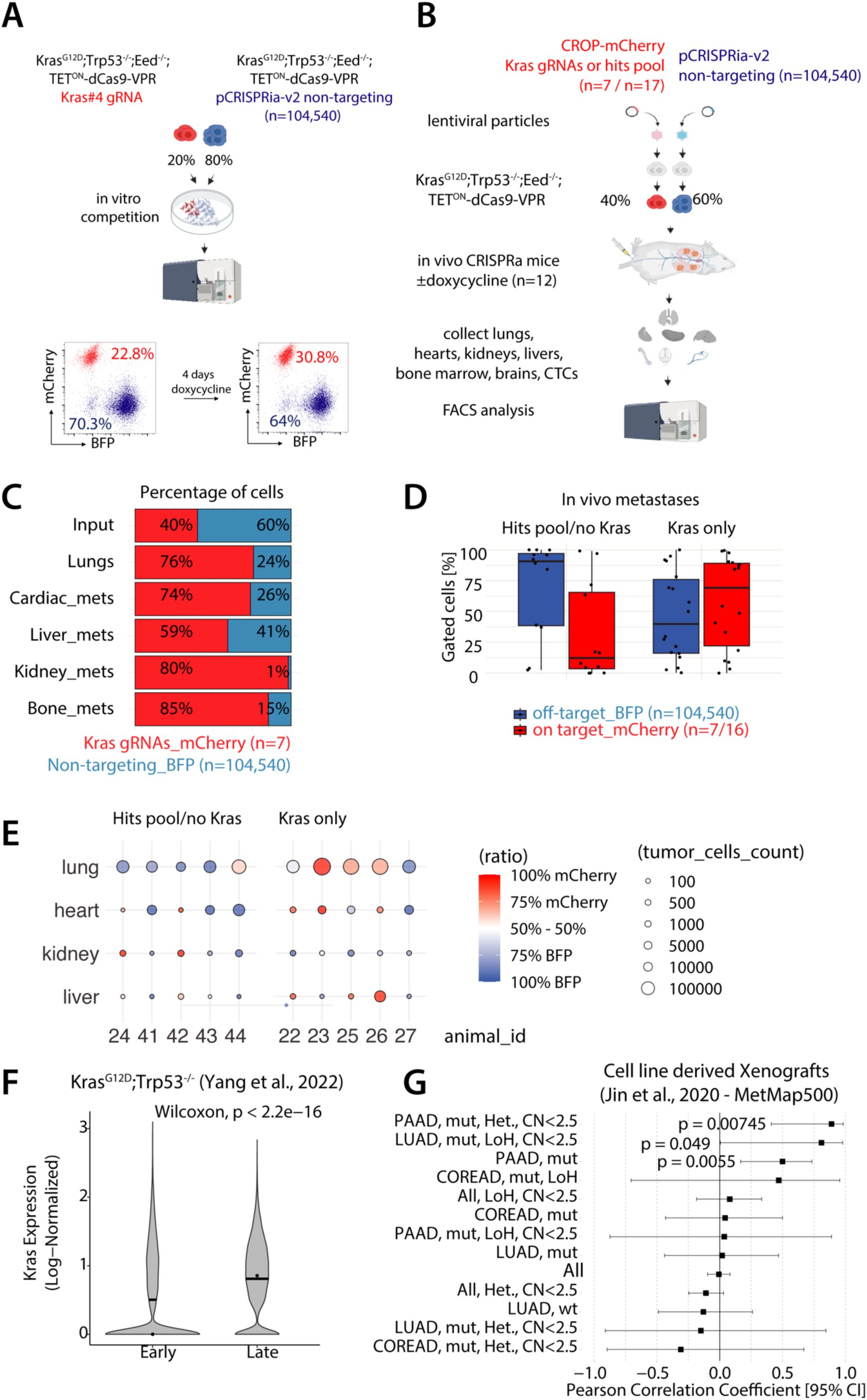
*Kras* transcriptional amplification by CROP-seq CRISPRa fuels lung cancer growth and metastasis **A)** Schematics of the cell competition experiments. Left, a representative FACS plot showing the dynamics of KPE;dCas9-VPR cells propagated under doxycycline and modified with either the CROP-seq-mCherry-gRNAKras#4 or pCRISPRia-v2-BFP non-targeting gRNAs. **B)** Schematics of the *in vivo* cell competition experiment between Kras-targeting or non-targeting pools. **C)** Stacked bar plot summarizing the multi-organ grafting competition experiment. FACS analysis of lungs, livers and kidneys (n=11), hearts (n=10) and bone marrows (n=7). **D)** Box plot comparing the distribution of percentages for mCherry and BFP expressing tumor cells of metastases from pro-metastatic pooled sgRNA or KRAS-only sgRNA overexpression. **E)** Dot plot illustrating the distribution of tumor cells within various organs for different experimental conditions. Each dot represents an individual sample with its size reflecting the number of tumor cells. The color of the dots represents the ratio of mCherry to BFP expression, with a gradient from blue (100% BFP) to red (100% mCherry). The x-axis shows the organ type, and the y-axis represents the animal ID. **F)** Violin plot of *Kras* normalized expression in single cells from autochthonous KP-tracer mice grouped according to disease stage. Data are from Yang et al., 2022. **G)** Forest plot of the Cox proportional hazards model illustrating the effect of KRAS expression on the metastatic patterns of CCLE cell lines from the MetMap500 (Jin et al., 2020). Pearson correlation coefficient is shown for the indicated sub-classes of cell lines divided according to cell line tissue of origin, KRAS status, copy number and zygosity.

We next set up a parallel competition assay *in vivo* between off-target gRNA bearing cells (BFP-labeled) against pooled gRNAs targeting either Kras or other random genes enriched in our primary screen with statistical significance but unclear functional role (**Fig.S3A-D**), which may be due to lower functionality, on-target activity, or off-target stochastic clonal expansion. *Kras* transcriptional amplification over remained significant whereas the other targeting gRNAs largely lost the competition to the control cells (**Fig. 3D- E)**. Some gRNAs showed dominant enrichment over others (**Fig. S3B-D**), yet their functional weight is lower than *Kras* and potentially independent from their target gene. The finding that only *Kras* but not other previously validated oncogenes such as Vav1, Cxcr4 and L1cam significantly accelerated progression under physiologically-relevant dosage amplification was rather surprising in that Kras^G12D^ is the genetic driver event used to initiate tumorigenesis in our autochthonous model ^20^. Moreover, in line with the standard mode of Kras-driven tumorigenic in autochthonous pancreatic cancer^26^, both the wild-type and mutant *Kras* alleles are genomically amplified also in KPE cells^28^. Biochemically, we confirmed that most *Kras* targeting gRNAs drove mRNA amplification of both wild-type and mutant allele, and therefore including but not limited to the oncogenic Kras protein (**Fig. S3E-F)**. In line with expectations for an endogenous system like CRISPRa targeting a highly expressed gene, we note that *Kras* amplification was mild and limited to two-to-four folds over basal levels in KPE (**Fig.S3D-E)**.

In a reanalysis of single-cell RNA sequencing and lineage-tracing data by Yang et al. ^18^, we next found that increased *Kras* transcriptional dosage is significantly linked to late-stage tumorigenesis and metastatic phenotypes in the autochthonous KP Tracer model (**Fig. 3F**). Notably, *Kras* detection rates, serving as a proxy for transcriptional levels, were markedly elevated in mesenchymal and metastatic cells, which dominate the late-stage clusters in an unperturbed metastatic setting (**Fig. S3G-H**). Next, we aimed to assess the role of *Kras* dosage in a human xenograft setting. Our analyses of data from the DepMap and MetMap500 projects demonstrated that transcriptional dosage of oncogenic *KRAS* correlates with metastatic potential and tissue colonization of human lung and pancreatic cancer cell lines across five organs upon xenografting (**Fig. 3G**). Notably, this effect was predominantly driven by the mutant allele, as allelic imbalance with the wild-type *KRAS* allele emerged as a tissue-specific feature (**Fig. 3G**), and the *KRAS* dosage effect was abrogated when non-mutant cell lines were analyzed (**Fig. S3I-M**).

In summary, transcriptional amplification of oncogenic KRAS drives progression and metastasis in KRAS^G12D^-driven cancers across multiple experimental models, underscoring the broad relevance of KRAS dosage in driving metastatic potential.

### *KRAS* transcriptional levels are linked to poor survival in primary KRAS-mutant driven cancers

If increasing *KRAS* transcriptional dosage during progression has functional relevance in human cancers, there should be correlative evidence in primary and metastatic datasets. Consistently, significantly shorter survival was associated with *KRAS* expression in patients with primary LUAD (TCGA) adjusted for age and KRAS mutational status. LUAD patients divided in groups according to *KRAS* transcriptional dosage, after having defined its impact across expression bins (methods), also support that high KRAS dosage holds a mild but significant poor prognosis (**Fig. 4** and **Fig. S4B)**. Poor prognosis and KRAS transcriptional dosage remained significantly associated even when the Cox model was extended to metastatic META-PRISM (PRISM) and Hartwig Medical Foundation (HMF) cohorts (4)(31), and other cancer types in which oncogenic KRAS is either the undisputed genetic driver (e.g. PAAD) or a well-established genetic driver of progression (COREAD=COAD + READ; **Fig. S4A**). Of note, in primary TCGA and metastatic HMF LUAD cohorts the RAS84 gene signature ^13^ that focuses on KRAS activity rather than transcriptional levels performs comparably to *KRAS* gene alone (**Fig. S4E-G)**. This is consistent with a model in which *KRAS* transcriptional dosage is an early driver of progression and metastasis.

**Figure 4.**
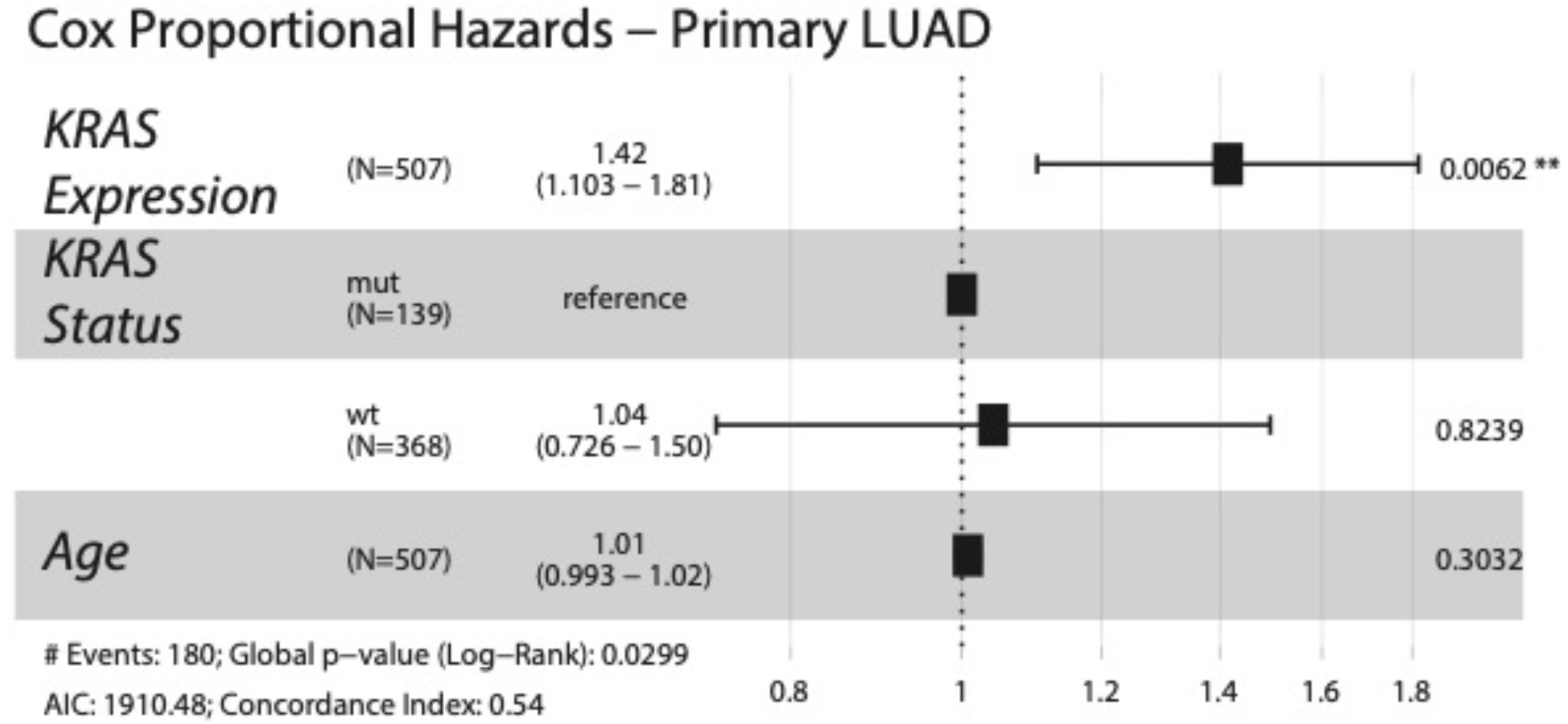
KRAS transcriptional levels are linked to poor survival in primary lung adenocarcinoma. Forest plot of the Cox proportional hazards model illustrating the effect of KRAS expression on survival in TCGA-LUAD. KRAS expression is adjusted for KRAS mutation status and age (see methods). P-value and hazard ratio of a covariate is indicated in the right and middle of a row respectively. Asterisks denotes significance. Global p-value is indicated at the bottom.

### Kras-driven cells interact with a sterile inflammatory microenvironment during tumor progression and with the activated TGFB pathway during metastasis

To assess the fidelity of our transplantation KPE model compared to the autochthonous mouse KP model, we exploited the parallel mRNA information obtained in our screening procedure (**Fig. S5A-B**). Unbiased clustering analysis showed that KPE extracted from orthotopic lungs adopt a homogeneous transcriptional state, whereas metastases display broader inter-sample variability (**Fig. S5B**). Gene set enrichment analysis (GSEA) shows that orthotopic and metastatic KPE display signatures associated with a broad set of the KP mouse cell states ^18^. A transition from a state resembling endoderm-and gastric-like, pre-EMT, and high plasticity cell states converge onto the full MES1/2 states ^18^ (**Fig. S5C)**, supporting the pathophysiological relevance of our model.

To exploit our model for the retrospective assessment of the oncogenic pathways whose parallel activation cooperates with Kras in driving LUAD progression in the mouse, we performed Ingenuity pathway analysis (IPA) of ex vivo KPE transcriptomic profiling derived from either the lungs or metastatic sites. This revealed that KPE cells experience inflammatory signaling via the JAK/STAT and NF-kB pathways, which was prominently sustained in the lung and retained, albeit to a lower extent, in the metastatic sites (**Fig. 5A)**. Differential IPA between metastatic and orthotopic tumor-derived cells indicates that KPE cells acquire a signature of TGFB signaling exposure once they left their orthotopic niche (**Fig. 5A)**. To validate the computational analyses, we reversely engineered the lung tumor microenvironment and *in vitro* subjected KPE cells to the cocktail of pro-inflammatory cytokines inferred from IPA *ex vivo*. We used Il33 and Ncam1 as a proxy of sterile inflammation-and TGFB-driven cell state changes, respectively (**Fig. 5B)**, and combined Interferon gamma, TNFa, Il1B and OSM as pathophysiologically relevant lung TME mimic (**Fig. 5A)**. RT-qPCR follow-up validated both the induction of Il33 by sterile inflammation and the shift towards Csf1 and Ncam1 expression by the addition of TGFB1 (**Fig. 5C)**. As an independent approach, we used a high-content analysis (HCA) screen for Ncam1 expression (**Fig. 5D)**, which aimed at the unbiased discovery of external signaling driving Ncam1 expression and potentially mirroring a metastatic niche signaling. This screen nominated TGFB1 as the sole external factor leading to Ncam1 activation KPE cells exposed to inflammatory signaling (**Fig. 5E-F)**, and this *in vitro* setting as driver of biomarkers observed *in vivo*.

**Figure 5.**
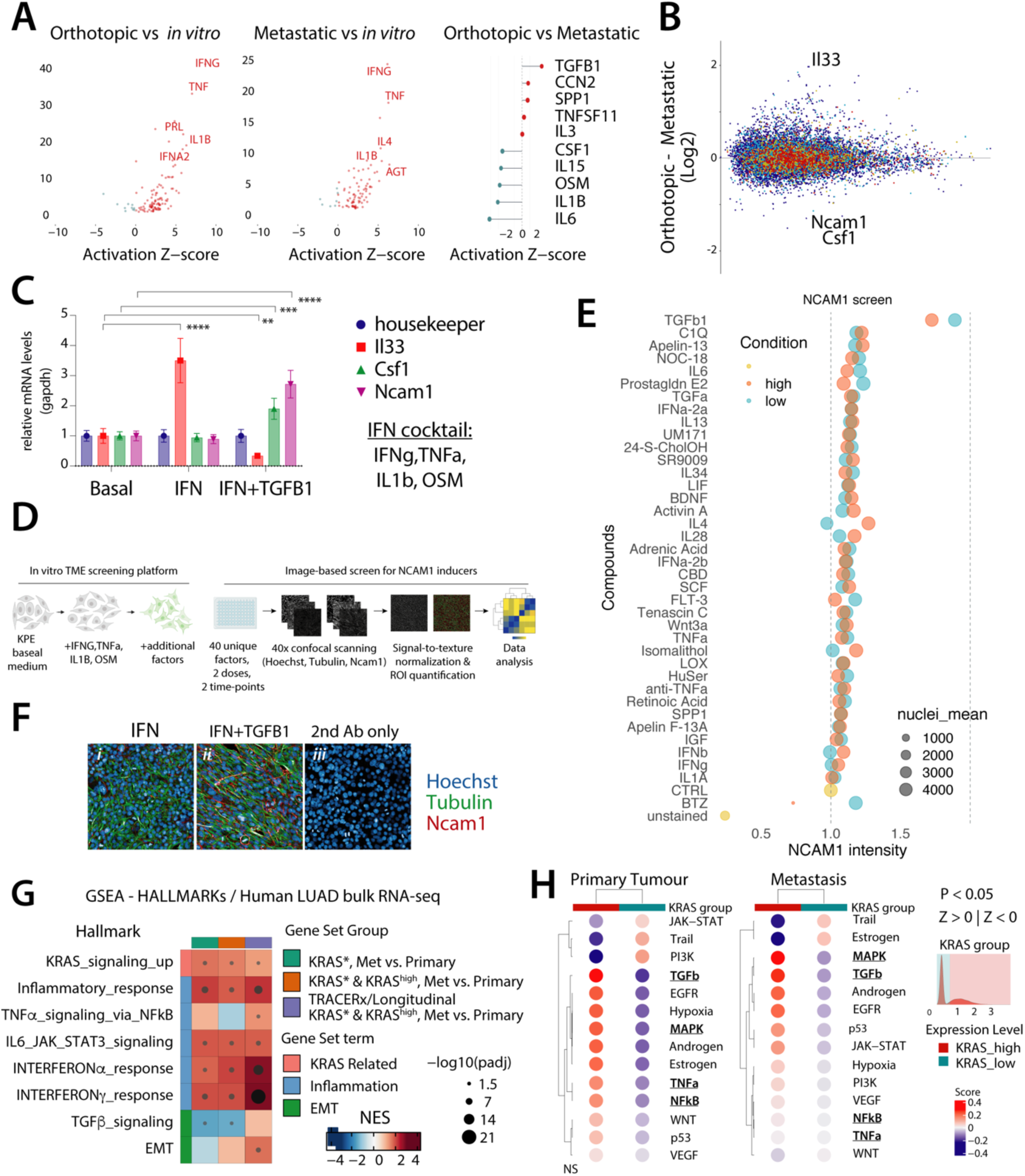
Kras-driven cells interact with a sterile inflammatory microenvironment during tumor progression and with the activated TGFB pathway at a metastatic site. **A)** Upstream regulators analysis for the indicated comparisons of orthotopic (n=11), metastatic (n=45) and in vitro (n=17) samples. Significant differentially-regulated genes were identified using LIMMA (padj<0.05, FC±0.5), Top ten regulators were identified using Ingenuity Pathway Analysis. **B)** MA plot showing differential expression between metastases-(n=45) and lung-or (n=17)-tumor derived lines. **C)** Bar plot showing RT-qPCR data for the indicated genes and conditions. P values were calculated by two-way ANOVA and Dunnett post-hoc test (only significant comparisons were shown; ****, P ≤ 0.0001; ***, P ≤ 0.001, **, P ≤ 0.01). **D)** Workflow of the pro-metastatic upstream regulators screening: lung orthotopic tumor TME signaling was inferred from (A) and were Ncam1 from (E) is used as biomarker for metastatic TME signaling. **E)** Scatter showing regulators of Ncam1 protein expression as proxy for metastatic TME in KPE cells exposed to inflammatory mediators from (A). **F)** Immunohistochemical staining for the indicated markers in KPE in inflammatory medium (i), TGFB1+ 10ng/ml (ii). A no secondary antibody control is also shown (iii). **G)** Heatmap showing GSEA normalized enrichment scores and adjusted p-value for the indicated gene sets. Asterisks denote p-value level: *, p < 0.05; **, p < 0.01; ***, p < 0.001. Patients are divided according to their known *KRAS* mutation status and disease stage (legend). The broad class of the gene sets is also shown in legend. **H)** Dotplot heatmap showing activity scores (presented as z-score) of PROGENy pathways (left) in human primary and metastatic cancer cells stratified by KRAS expression levels (Laughney, AM et al n=17 patients, 35,985 total cells, 1,731 aneuploid cells).

To corroborate our experimental findings in distinct models, we ran PROGENy pathway analysis on the single-cell RNA-seq data by Yang et al. and found that the JAK/STAT pathway activation is particularly connected with high KRAS dosage, whereas the TGFB activity increases during progression (**Fig. S5D**). In the human setting, we next applied gene set enrichment analysis to an ensemble of primary and metastatic bulk RNA-seq profiles from a total of 1467 LUAD patients. This includes molecular datasets of primary (TCGA, n=516) or metastatic (PRISM, n=165 and HMF, n=284) tumors but also longitudinal multisampling by TRACERx data (n=502, longitudinal n=80 from 17 patients), which encompass the full spectrum of tumor evolution, from primary to metastatic stages ^34^. This analysis was well-powered to test the association between a high *KRAS* transcriptional dosage and the activation of pathways connected to inflammation and fibrosis during LUAD progression. GSEA using the classic Hallmarks gene sets selected for KRAS activity, inflammation and EMT, revealed that indeed inflammation plays an essential role in oncogenic KRAS-driven LUAD progression and metastasis (**Fig. 5G**). As complementary computational approach, we used PROGENy, which is designed specifically on human cancer data to detect pathway activation. Both analyses converged on JAK/STAT pathway connection with a high *KRAS* dosage in KRAS mutant during metastatic progression, including in the direct TRACERx longitudinal cohort (**Fig. S5D**). The confirmatory finding that high KRAS transcriptional dosage strongly correlates with the MAPK pathway activation supports the robustness of the analysis despite the bulk nature of the transcriptomics (**Fig. S5E**). An interesting divergence between the mouse experimental and the human descriptive datasets is the opposite behavior of the TGFB pathway, which appears to suggest that the metastatic stage in the models is distinct from that measured post-mortem in humans (see discussion).

Still, bulk RNA-seq analyses remain potentially confounded by the microenvironment, even if correcting for transcriptional purity is possible. To confirm that KRAS transcriptional levels converge with specific pathways during progression and metastasis in human cancer cells, we analyzed single cell datasets, which can be restricted to aneuploid cells. To this end, we used two independent reference datasets featuring either a limited but matched primary metastatic dataset in western patients for KRAS LUAD ^35^ and single cells from or a deeper non-small cell lung cancer dataset in the Chinese population ^36^. After filtering for aneuploid cells based on RNA-seq inference of copy-number aberrations and divided cell in two groups according to *KRAS* transcriptional dosage (**S5F-H)**, PROGENy confirmed that high *KRAS* LUAD cells, whether primary or metastatic, feature the simultaneous activation of the NF-kB and TGFB pathways when KRAS dosage is highest, along other pathways, including MAPK (**Fig. 5H**). From our analyses, whether mouse or human, bulk or single cell, the expected correlation between KRAS transcriptional dosage and MAPK pathway activity occurred as expected, whereas inflammatory or TGFB activity appeared to be dependent also on the early or late stage of progression, respectively.

Together, our data support a model in which high oncogenic KRAS dosage collaborates with sterile inflammation to drive progression and metastasis consistently across KPE, KP, and human data, and suggest that the TGFB pathway activation holds stage-specific functions.

### Kras transcriptional amplification operates within the context of a pre-existing chromatin configuration

The KRAS-driven signaling cascade drives MEK-ERK activation and results in a transcriptional output, which in turn sets the stage for gene expression patterns and acquisition of defined cell states ^37^. To obtain a mechanistical insight into how high KRAS transcriptional dosage drives tumor progression, we combined our *in vivo* model of KRAS transcriptional amplification by CRISPRa and our *in vitro* reverse engineering of LUAD microenvironment. Next, we conducted the Assay for Transposase-Accessible Chromatin using sequencing (ATAC-seq) in all of the above-describe conditions, as probing chromatin accessibility via ATAC-seq can potentially uncover both current, memory and potential cell states (**Fig. 6A** and **Fig. S6A)**. Overlap between independently called peaks showed that KPE cells share a large fraction of open chromatin, which is accessible independently from the growth conditions (*in vitro* and *in vivo*), external signaling (basal, pro-inflammatory/fibrotic *in vitro*, or TME *in vivo*), as well as KRAS transcriptional dosage (**Fig. 6B and fig S6A-E)**. We also observed that each condition is associated with selective patterns of qualitative (i.e. peaks) and quantitative (**Fig. 6B** inset, and **Fig. S6B-E)**, indicating that KRAS transcriptional dosage integrates its activity within the context of a pre-existing chromatin configuration.

**Figure 6.**
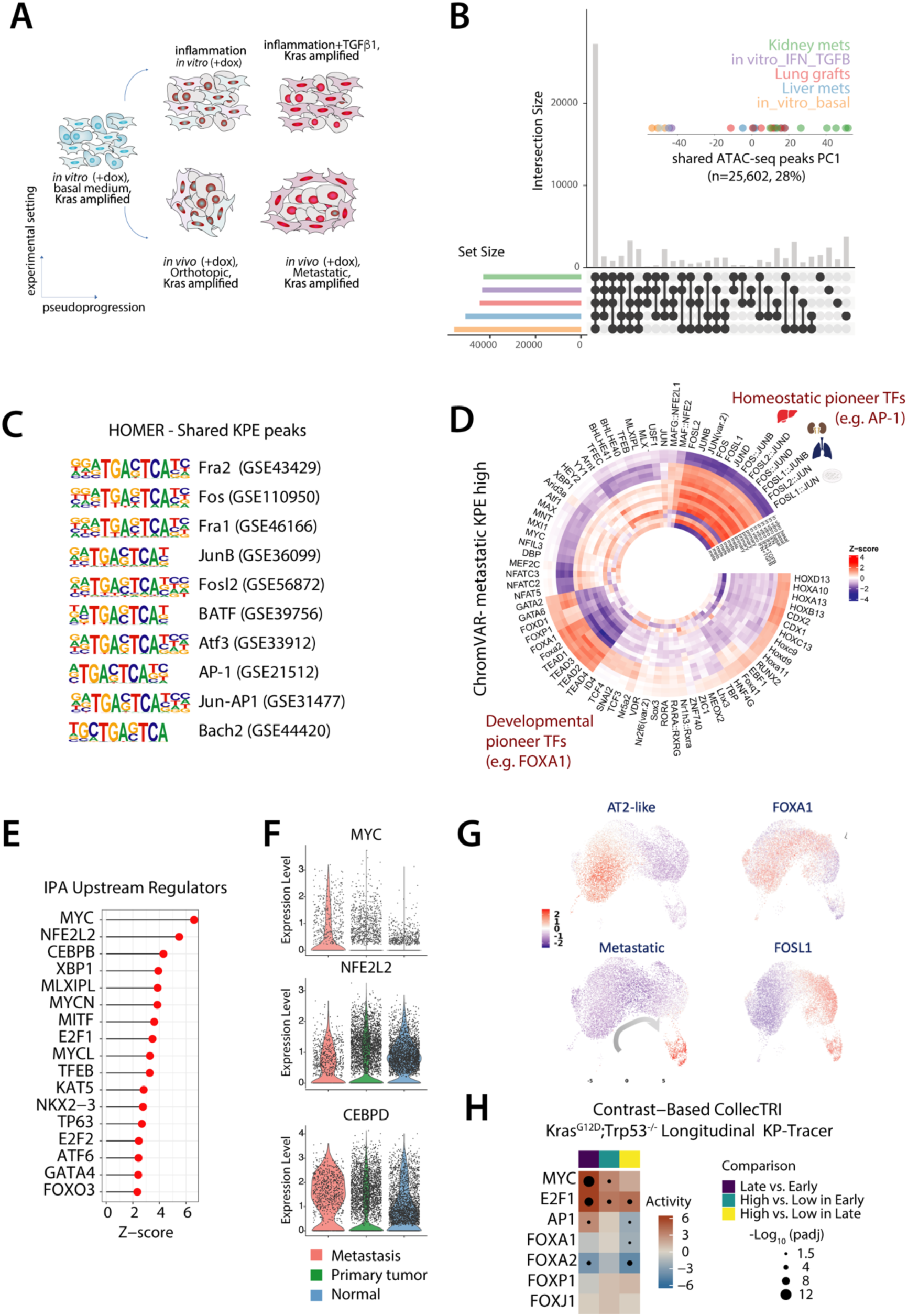
Kras transcriptional amplification operates via transcription factor dynamics at pre-existing open chromatin **A)** Schematic representation of the experimental conditions included in the ATAC-seq profiling. **B)** Upset plot showing the overlap between ATAC-seq profiles. Inset shows first principal component. **C)** Top 10 motifs predicted by HOMER TFBS enrichment analysis of overlapping ATAC- peaks from (B). **D)** Circular plot showing the ChromVar of ATAC-seq peaks specifically remodelled in vivo. Differential Z-scores deviation for the top TFBS motifs between each condition and basal in vitro is shown. **E)** Lollipop plot showing IPA upstream transcriptional regulators analysis of genes annotated near ATAC-seq peaks from (C). **F)** Violin plot showing expression levels of the indicated genes in single cells in human normal lung, primary and metastatic cancer cells (Laughney, AM et al n=17 patients, 35,985 total cells). **G)** UMAP representation of single-cell ATAC-seq module scores for the indicated pioneer factors in KP cancer cells from LaFave et al. The AT2-like (i.e. candidate LUAD cell of origin) and metastatic cell states and arrow indicate the spatiotemporal evolution of the mouse autochthonous LUAD. **H)** Heatmap showing activity scores of CollecTRI transcription factor activity inference in in single cell clusters associated with progression stages and separated based on Kras expression as in Fig. 2F from the autochthonous KP-tracer model as defined by scRNA-seq and lineage-tracing (Yang et al., 2022). Asterisks denote p-value level: *, p < 0.05; **, p < 0.01; ***, p < 0.001.

To infer the transcription factors that prime chromatin for KRAS signal transduction in KPE, we next ran HOMER associated on KPE shared open chromatin. This reveled that the main transcriptional factors marking this open chromatin landscape in KPE are the individual factors composing the main configurations of the AP-1 dimer (**Fig. 6C)**. Next, we probed differentially accessible regions (DAR) that are significantly open in metastasis-derived KPE compared to basal (**Fig. 6B** and **S6C)** using ChromVAR and found that AP-1 featured the most relevant driver of such accessibility, indicating that KPE progression is associated with AP-1 hyperactivation. The samples that were subjected to *in vitro* stimulation with pro-inflammatory and/or fibrotic factors, offered support to this analysis, as AP-1 appeared to be activated specifically by pro-inflammatory signaling (**Fig. 6D**), which follows consensus ^38^.

To gain insights into the potential transcriptional regulators of the KPE accessible chromatin, we annotated genes nearing the shared peaks and performed IPA upstream regulators analysis. This uncovered that MYC/E2F, NRF2 (i.e. NFE2L2) and CEBP factors are the main predicted regulators downstream oncogenic KRAS signaling in KPE (**Fig. 6E)**. Of note, all of these factors are much more expressed in LUAD single cells from primary and metastatic sites, if compared to resident cells in human lungs, and MYC and CEBPD are particularly enriched at metastatic presentation (**Fig. 6F)**.

Together, the data support a model in which signal transduction by KRAS promotes the activity of highly expressed transcription factors within the context of pre-existing open chromatin.

### Chromatin remodeling during lung cancer progression *in vivo* reveals a pioneer transcription factor transition

Tumor cells face the need to adapt to different environments and execute diverse transcriptional programs, but many of the pathways and factors herein identified are largely connected to proliferation. In the ChromVAR analysis, we noted a marked shift in accessibility of AP-1 associated TFBS, whereas TFBS associated with GATA, TEAD, and FOX transcription factors followed the opposite trend as AP-1 (**Fig. 6D**). FOX factors are canonical developmental pioneer transcription factors, capable of accessing their cognate motif even when embedded into an intact nucleosome ^39^. To gain pathophysiological relevance for this pioneer factors dynamics, we exploited an ATAC-seq single-cell profile previously generated from ex vivo autochthonous KP LUAD and metastases ^19^. Importantly, we confirmed that data generated in our KPE model follow the same trend as within an intact mouse model, with FOX TFBS being progressively closed as cell state progression towards metastasis is plotted. In contrast, but perfectly aligned with our predictions, AP-1 TFBS are consistently more accessible at late stages of the tumorigenic process and metastasis (**Fig. 6G**).

We next wished to corroborate such transcription factor dynamics in the autochthonous KP tracer progression ^18^. To this end, we used CollecTRI to infer the TF activity inferred in human cancers in single-cell longitudinal transcriptomic profiles from early and late/metastatic mouse LUAD cell states. This analysis shows that MYC, E2F and AP-1 targets are significantly associated with Kras-high expression and enriched during progression, whereas and FOXA1, FOXA2, FOXP1 follow the opposite trend, with FOXA2 targets being particularly (**Fig 6H**). This analysis confirmed that AP-1 activity appears to correlate with progression in human LUAD as well.

Since MYC, NRF2, CEBPB, and E2F are DNA binding factors expected to bind accessible cognate TFBS whereas both FOX and AP-1 factors are considered pioneer transcriptional regulators capable of binding nucleosomal DNA, the observed switch between FOX and AP-1 during tumor progression supports a model in which high KRAS activity cooperates with pioneer transcription factor dynamics to potentially dictate cell states.

### Kras dosage dictates Kras-driven cells states and explains responses to KRAS Inhibition

Having connected a high Kras dosage and tumor progression with external signaling and pioneer transcription factors dynamics *in vivo* (**Fig. 6D-G**), we next wished to dissect the net contribution of each component over KPE cell state dynamics. To this end, we conducted both RNA-seq and ATAC-seq on KPE cells in which individual components were perturbed. On the one hand, we exploited the ability of inflammatory cytokines and TGFB1 to impart a transcriptional program that amplifies AP-1 activity without impacting on Fox transcription factors (**Fig. 6D)**. On the other hand, we deleted the most expressed developmental pioneer factors in KPE, *Foxa1* and *Foxp1*, using CRISPR-based genetic knock-out. Finally, we transcriptionally modulated Kras dosage by generating an additional set of KPE lines carrying a repressive dCas9-MeCP2, directed to the Kras locus via the same gRNAs for CRISPRa. Strikingly, both ATAC-seq and RNA-seq showed concordant variation across the two major PCA components and the individual replica experiments remained close, indicating that both chromatin and transcription lie on a single axis of variation dependent on *Kras* dosage (**Fig. 7A-B**). As the second axis of variation successfully resolves the phenotypic impact of the pro-inflammatory phenotype and the loss of pioneer factors’ activity, this indicates that, in our model, KRAS dosage appears to dominate transcriptional and chromatin accessibility variation (i.e. PC1). To investigate how *Kras* dosage modulation imparts specific cell states in KPE globally, we defined the upstream regulators of differentially expressed genes using IPA. In this analysis, we independently calculated the upstream regulators in KPE cells in which only Kras dosage was modulated (KPE), or Kras dosage was modulated in presence of inflammation and TGFB1 (KPE+IFN+TGFB) or absence of developmental pioneer transcription factors (KPE;Foxa1^-/-^;Foxp1^-/-^;+IFN+TGFB). After that, we clustered the results and observed that a high Kras dosage holds a dominant effect over transcriptional networks associated with either a proliferative cell state (e.g. Myc, E2f, Srf), whereas a low Kras dosage promotes the upregulation of transcriptional programs connected to stress responses (Atf4, Nrf2, Stat3; **Fig. 7C)**. Of note, low Kras dosage also induced a transcriptional program connected to Foxa1. Hence, we performed an immunoblot to probe each individual component and discovered that Kras dosage antagonistically modulates the AP-1 and Foxa1/Foxp1 protein levels (**Fig. 7D-E)**. GSEA confirmed a dominant role for Kras dosage over the other components when focusing on the regulation of genes residing in chromatin remodeled during metastasis (**Fig.S7A)**.

**Figure 7.**
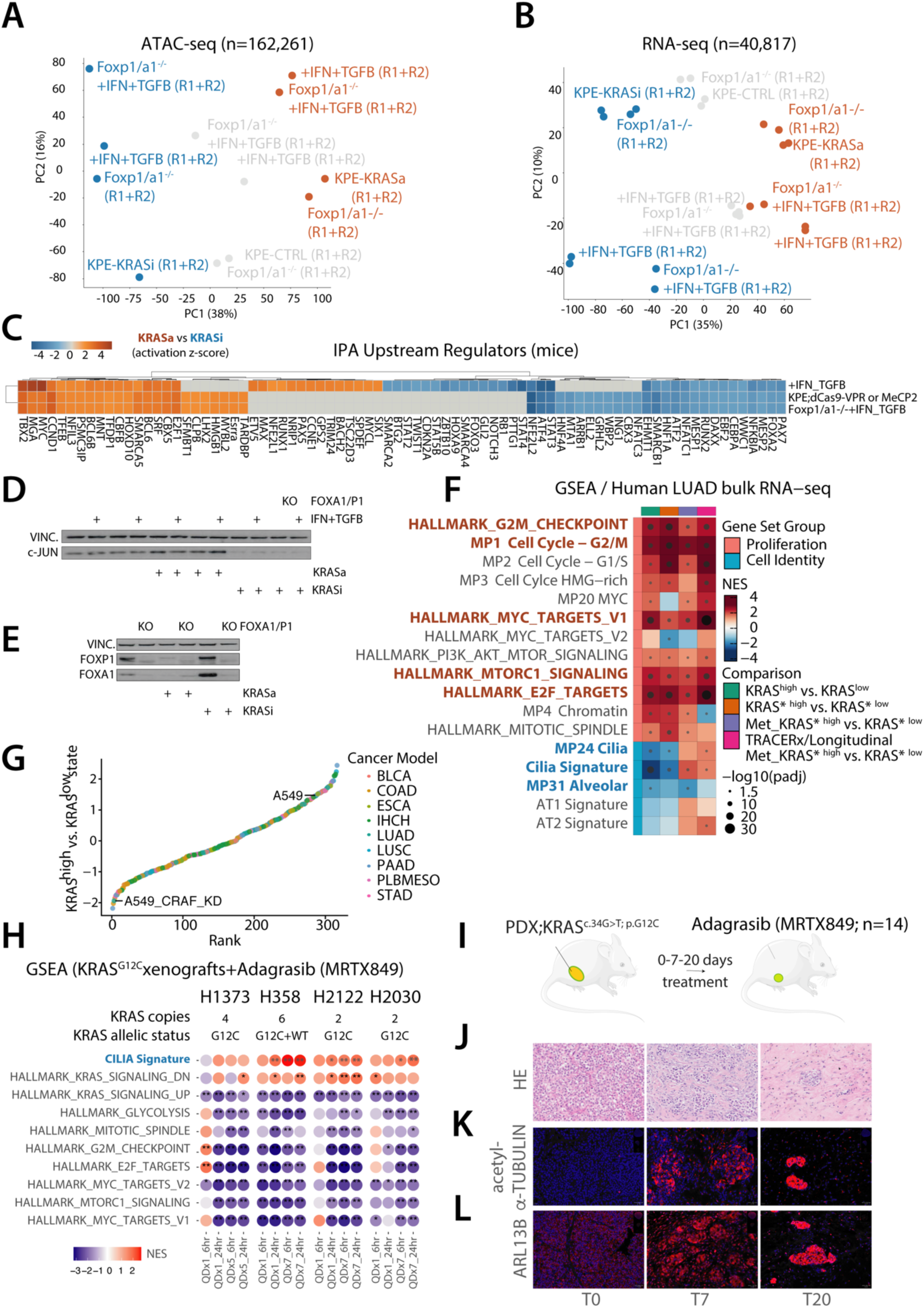
Kras dosage dictates Kras-driven LUAD cells states. **A)** PCA of a broad set of accessible peaks from KPE cultures with the indicated genotypes and treatments. Dot are colored according to their KRAS dosage, determined by the presence of gRNAs targeting the Kras locus, and doxycycline inducing either an activating dCas9-VPR (orange) or repressive gCas9-MeCP2 (blue). **B)** PCA of a broad set of mRNAs from KPE cultures with the same setting and color code as in (A). **C)** Upstream regulator analysis by IPA for the indicated comparison on the genes passing the FC threshold (FC±0.5, padj<0.05). **D-E**) Western blotting of the indicated antibodies in lysates from KPE cultures with the indicated genotypes and treatments. Note the dominant role for KRAS dosage in c-JUN and FOX factors protein levels. Foxa1 and Foxp1 KO controls as immunoblot specificity and sensitivity are shown. **F)** Heatmap showing GSEA scores (presented as normalized enrichment scores, NES) of the indicated pan-cancer single-cell consensus meta-programs and selected hallmarks in KRAS high vs. low contrast setting (see methods) in primary and metastatic bulk RNA-seq profiles. Patients are divided according to their known KRAS mutation status (“*” legend). Filled circles denote Benjamini-Hochberg adjusted p-value level: *, p < 0.05; **, p < 0.01; ***, p < 0.001. **G)** Heatmap showing GSEA scores (presented as normalized enrichment scores, NES) of the Cilia signature associated with KRAS dosage in LUAD in (F) and hallmark signature pathways differentially regulated in at least one Cell line derived xenograft (CDX) 24 hours following oral administration of a single 100 mg/kg MRTX849 dose compared with vehicle. NES shown in all models 6 or 24 hours after a single dose (QD × 1) or 5 (QD × 5) or 7 (QD × 7) days dosing. Data were reanalyzed from Hallin et al. **H)** Schematic of a LUAD KRAS^G12C^ patient-derived xenograft (PDX) experiment to test the phenotypic markers associated with minimal residual disease following continuous administration of MRTX849. **I)** Representative hematoxylin-Eosin (HE) staining of PDX subcutaneous tumors treated with Captisol (left), and 100 mg/kg MRTX849 QD × 7 or QD × 20 (center, right, respectively). **J-K**) Representative immunohistochemical (IHC) images of PDX tumors collected at indicated timepoints, stained for acetyl-Tubulin (j, red) or ARL13B (k, red) and counterstained with DAPI (blue). Scale bars = 20 μm.

To gather evidence that *KRAS* dosage associates LUAD progression and metastasis to gene expression via distinct cell state programs, we next exploited the compendium of human pan-cancer cell states recently generated by non-negative matrix factorization of single-cell RNA-seq data ^40^, along with well-established hallmark gene sets. GSEA contrasting patients’ biopsies with high or low KRAS transcriptional dosage in primary LUAD (TCGA, TRACERx), and metastatic LUAD (PRISM, HMF, TRACERx), showed that a high KRAS dosage positively correlates with proliferation cell states, MYC and secretory activity and a EMT state independently of the KRAS gene mutation status and LUAD progression (**Fig. 7F**). In contrast, a low KRAS dosage is associated with cell differentiation, with Alveolar, AT1 and Cilia cell states marked by statistical significance in the primary LUAD but not metastasis (**Fig. 7F**). These states were the only ones recurrently associated with KRAS transcriptional dosage among all the pancancer single cell states and several gene sets associated with cancer hallmarks (**Fig. S7B)**. Strikingly, unlike the AT1 state that appears to be strongly connected to the *KRAS* dosage independently of the mutational state and LUAD stage, the “ciliated-like” showed a stage-dependent switch in the mutant, raising the intriguing possibility that this cell state may hold functional properties.

The use of single-cell RNA-seq and large number of biopsies from distinct biopsies support the theory that these cell states reflect a cell intrinsic cancer cell property directly regulated by KRAS signaling. Consistent with this hypothesis, all solid tumor lines from the Cancer Dependency Map portal (DepMap) can be categorized in KRAS-high, KRAS-low states and non-classified (**Fig. S7C**). Albeit we did not find evidence of “ciliated-like” states in LUAD cell lines, we discovered that that the quasi-mesenchymal A549 LUAD cell line resembles KRAS-high patients and switches to a KRAS-low phenotype when RAF1 is knocked-down (**Fig. 7G**). Thus, the KRAS-high and KRAS-low ends of the transcriptional spectrum we discovered in LUAD patients depend on *KRAS* dosage via the canonical RAS/RAF signaling.

Direct KRAS inhibition (KRASi) disconnects KRAS dosage from its downstream activity and leads to downregulation of KRAS, E2F and MYC signatures in xenografts^41^. In the KP mouse model, AT1 differentiation stabilizes KP cells in response to KRASi^17^. Thus, we hypothesized that direct KRAS inhibition would directly target the KRAS-high proliferation cell state and drive a KRAS low-like phenotype at minimal residual disease level. To test this, we run a differential GSEA in four cell line derived xenografts (CDX) treated *in vivo* with the clinically-approved covalent KRASG12C inhibitor MRTX849 (a.k.a. Adagrasib). This analysis also confirmed our previous finding that KRAS drives EMT^20^ and supported the herein describe connection between KRAS dosage and TGFB activity (**Fig. S7B**). Importantly, this analysis also showed that KRASi induces the Ciliated-like state (**Fig. 7G**), which we had found significantly associated with primary LUAD in patients cohort but not in established cell lines, suggesting that such state occurs under *in vivo* (**Fig. 7F**). Of note, the conserved biomarker of ciliated lung epithelial cells is FOXJ1, which is significantly inversely correlated with MYC, E2F1, NF-kB and AP-1 (**Fig. 6H** and **S6F**).

To prospectively test our hypothesis, we established a KRASG12C LUAD patient-derived xenograft (PDX) model, and treated F4-generation mice (n=22) with MRTX849 or vehicle. Prolonged KRASi (up to 67 days) induced a marked reduction in tumor volume, which was reversed upon drug removal indicating that KRAS inhibition induces the establishment of MRD that holds intact potential to restart tumor growth (**Fig. S7C-D**). Response to KRASi was accompanied early on by a substantial shift in tumor phenotypes from epithelial and proliferative-like to fibrotic-like (**Fig. 7H**). IHC highlighted a high proliferative index in the pre-treatment lesions as illustrated by cellularity, Ki-67 and phospho-H3 positivity (**Fig. S7E** and data not shown), suggesting high MYC/E2F activity in agreement with our data and predictions based on literature. Most importantly, residual tumor cells, which hold intact potential to restart tumor growth upon KRASi withdrawal (**Fig.S7C)**, showed substantial acetyl-alpha-TUBULIN and ARL13B positivity, both of which are well-established markers of ciliated cells (**Fig. 7I-K**). Hence, data in CDX and PDX models support the significance of a previously not captured Cilia-like cell state in both tumor evolution and response to targeted therapy.

Taken together, our data nominate oncogenic KRAS dosage as the key driver tumor evolution shaping cell states during mouse and human LUAD progression and metastasis as well as in response to direct KRAS inhibition.

## Discussion

Successful first-line cancer treatments renders metastasis an uprising societal challenge. Therefore, understanding the mechanistic basis of metastasis is crucial. We identify oncogenic KRAS dosage as a key driver of LUAD progression by integrating external signaling and pioneer transcription factor dynamics into distinct cellular states, which are key in driving tumor evolution and response to KRAS inhibitors ^8–11^.

Our results extend beyond the KPE model. Validation in KP and CDX models underlines that KRAS dosage influences lung adenocarcinoma progression in genetically and physiologically diverse systems. These results suggest that KPE offers unique insights into advanced cell states and that the KRAS dosage effect is translatable across models.

Although a single aberration is typically insufficient to transform a human cell ^42^, KRAS mutations meet the established criteria for cancer drivers ^13^, particularly in LUAD or PDAC where they feature initiating genetic event. Here, we demonstrate that increasing the transcriptional dosage of oncogenic KRAS is sufficient to drive LUAD progression. This aligns well with previous observations in the autochthonous PDAC model ^26^, and our endogenous transcriptional modulation via CRISPRa is now able to disentangle the transcriptional dosage from copy number aberrations. Moreover, our in vivo transplantation LUAD model separates initiation from progression and metastasis. Both the KP model in Muller *et al.* and the KPE model used here rely on KRAS^G12D^, bear spontaneous KRAS genetic amplification and retain the wild-type allele. However, distinct KRAS mutations have unique functional properties ^43^, and role of the wild-type allele in KRAS mutant cancers on progression and response to targeted therapy is significant yet context-dependent^44–47^. Nevertheless, our data extend beyond the model employed since subclonal expansions within primary LUAD are linked to poor survival independently of KRAS ^48^. Expanding our approach to cross-validate findings across both immunocompromised (KPE, human xenografts) and immunocompetent (KP model) systems reinforces that KRAS dosage impacts cell states and metastatic dynamics broadly, with elements associated with KRAS dosage being also recapitulated in other KRAS-driven and even KRAS-wild-type cancers.

From a biomarker perspective, the observation that mild transcriptional up-regulation of KRAS promotes clonal expansion of orthotopic and metastatic lesions supports the speculation that copy number aberrations may stabilize the KRAS-driven subclonal expansion as an advantageous trait in a Darwinian sense therefore representing a fraction of the KRAS-high lesions. If confirmed, KRAS transcriptional levels would hold prognostic value. Hence, it is worth noting the marked accumulation of polyclonal cancer cells within the heart vasculature (**Fig. 1B**), specifically in the pulmonary vein and aorta, highlighting a potential ‘sewage-like’ effect. It is plausible to consider a digital RT-qPCR screen of pulmonary vein samples in clinical studies on pulmonary hypertension or prospectively in patients with KRAS-mutant tumors at the minimal residual disease stage. This approach could be superior to cell-free DNA detection in bronchoalveolar lavage, which is considered for LUAD screening but is confounded by the presence of KRAS mutations in pre-transformed tissues.

From a therapeutic perspective, we recently proposed that targeting oncogenic drivers along with driver cell states may be key to attaining profound responses ^7^. When KRAS transcriptional dosage dominates LUAD progression and cell states, future strategies should aim to target KRAS mutants and the KRAS high dosage cell state simultaneously. Our data in primary and metastatic patients’ cohorts suggest that the KRAS-high state relies on PI3K/mTOR and VEGFA activation and provide molecular support for the previously suggested combinatorial partners for KRAS inhibitors ^41^, possibly integrating anti-angiogenic therapies ^49^. Notably, lowering KRAS dosage drives chromatin and transcriptional remodeling, mimicking direct KRAS inhibition, which holds therapeutic potential, possibly in an adjuvant setting. The ‘ciliated’ phenotype observed in patient data (**Fig. 7** and **S7**) aligns with predictions from cell line models ^50,51^. It was recently shown that direct inhibition of KRAS leads to differentiation to AT1 in mice ^17^ and we show an additional Ciliated-like phenotype in PDX data upon KRASi and in primary human data with low-KRAS dosage (e.g., TCGA). Low KRAS dosage boosts Foxa1, Foxa2, FOXJ1, and stress response factors’ activity (**Fig. S7F**). Therapeutically, it will be particularly important to target this MDR as KRASi appears to be insufficient to phenocopy their genetic target inhibition

Oncogenic KRAS and FOX collaborate to define tumor phenotypic identity in both LUAD and PDAC models ^30,52–54^. The collaboration between KRAS-low dosage and FOX observed herein might hold dormancy-like potential in lung niches via differentiated cell states, which have distinct metabolic requirements. Surviving cells upon KRAS downregulation in PDAC fail to activate its hardwired anabolic glycolysis ^55^, yet do not show signs of (mitochondrial) stress upon OXPHOS inhibition ^56^. Whether these KRAS-low lesions may be sensitive to OXPHOS inhibitors is an exciting question stemming from this work. However, unlike the widely represented KRAS-high cell state, characterized by proliferative traits, the KRAS-low phenotype may depend more on primary and metastatic niches and will require precise characterization in patients on treatment.

Mechanistically, this study reveals how tumor cells with shared oncogenotypes, exposed to broad pathways, adapt to specific environments and execute diverse transcriptional programs. A high KRAS dosage collaborates with inflammation and fibrosis signaling to promote a FOX-to-AP-1 pioneer transcription factor transition. We present both *in vitro* and *in vivo* evidence supporting this key observation, which we further confirm in independent autochthonous models (**Fig. 6)**. Whether AP-1 qualifies as a pioneer transcription factor ^57,58^ or a stripe factor ^59^, its role in maintaining accessibility to various co-binding partners explains how tissue-agnostic homeostatic and regenerative programs may be integrated into a site-specific transcriptional output via the available accessory transcription factors. This way, the broad KRAS-ERK-MYC-AP1 signaling cascade adeptly integrates niche-directed transcription in cell type/state-specific outputs. Hence, our data connect the activity of broad pathways mediating cell-intrinsic signaling by oncogenic KRAS via ERK ^60,61^ and TME-driven NF-kB and TGFB, to context-dependent transcriptional outputs required for cell state stabilization in autochthonous and foreign niches.

In summary, the modulation of cell identity and functional properties defined in this study underscores oncogenic KRAS dosage as the *‘driver of drivers*’. Oncogenic KRAS functions in LUAD and PDAC models are similar in the context of different organs, and KRAS is a model oncogene, suggesting a broader applicability of our data to other RAS oncogenes, if not beyond. Technologically, our unbiased in vivo approach using CRISPR activation and fate mapping strategy is scalable, and a parallel approach utilizing CRISPR interference could similarly identify genes crucial for maintaining competitive advantage and be closer to therapeutic effects than CRISPR knock-out. Conceptually, our findings illustrate a bimodal evolutionary trajectory in LUAD, where both high and low dosages of oncogenic KRAS dictate the extremes of a spectrum covering all conceivable cell states within KRAS-driven solid tumors. Echoing the Paracelsian principle that *‘the dose makes the poison*’ our framework simplifies the complex continuum of states observed in patients through functional dimensionality reduction and highlights the crucial role of oncogenic KRAS dosage in determining the biological impact of KRAS on tumor heterogeneity, progression, and therapeutic responses.

## Material and Methods

### Animal experiments

All animal experiments were performed in compliance with the EU regulation. All the experiments but the PDX were in strict compliance with the German Animal Welfare Act and approved by the Regional Office for Health and Social Affairs Berlin (LAGeSo) under license number G0094/18. All PDX experiments strictly complied with the protocols approved by the University of Leuven Animal Care and Use ethical committee project number P123/2021 and by the ethical committee Research UZ/KU Leuven (S67758).

### Cell culture, primary cell isolation, transfection

NSCLC KPE cell lines were isolated from KrasG12D/+;Trp53-/-;EED-/- genetic background, as published in Serresi et al 2016 and infected with Luciferase to monitor their growth (hereby KPE-luc)(Cancer Cell) Primary and metastatic cancer cells were isolated using the gentleMACS™ Octo Dissociator and the respective kits specific for each tissue.

Cells were propagated in DMEM/F12 medium supplemented with 10% FBS, and 5% penicillin and streptomycin, 4g/ml of hydrocortisone (Sigma), 5 ng/ml murine EGF (Invitrogen), Insulin-Transferrin-Selenium mix/solution (GIBCO). Peripheral blood was collected into EDTA-coated microtubes, and Tumor circulating cells were isolated from red blood cell by hypotonic lysis (eBioscience™ 10X RBC Lysis Buffer) and analyzed at the FACS for mCherry expression.

A549, H1944, H2030, H3122 cell lines were described before^20^ and were cultured in RPMI medium with 10% FBS, and 5% penicillin and streptomycin at 37°C in a 5% CO2–95% air incubator.

KPE-luc transfection was conducted using Fugene HD transfection reagent (Promega) with doxycycline-inducible plasmids: PB-TRE-dCas9-VPR (Addgene #63800) and PB-TRE-dCas9-KRAB-MeCP2 (Addgene #122267). Following transfection, cells were cultured in growth medium supplemented with Hygromycin (100 µg/mL) to select for stable transfectants.

### Tissue clearing and lightsheet microscopy

Mice were anesthetized by intraperitoneal injection of 150 mg/kg ketamine and 10 mg/kg xylazine and transcardially perfused with PBS.Tissue clearing was performed as previously described with modifications (Susaki, E. A. et al. Advanced CUBIC protocols for whole-brain and whole-body clearing and imaging. Nat. Protoc. 10, 1709–1727 (2015).In short, tissue was transferred to CUBIC1 (25 wt% Urea, 25 wt% N,N,N′,N′-tetrakis(2-hydroxypropyl) ethylenediamine, 15 wt% Triton X-100) and incubated at 37 °C shaking. Every other day CUBIC1 solution was exchanged until tissue appeared transparent (2∼4 days). Afterwards, samples were washed for 1 day with PBS at RT, refractive index matched with EasyIndex (LifeCanvas Technologies) at 37 °C and imaged with the ZEISS Lightsheet Z.1, image analysis and video rendering, we used ZENblack v 3.1, Arivis Vision4D v3.5.1 (Arivis AG) and Imaris v9.8.0 (Oxford Instruments), respectively.

### Bioluminescence analysis and quantification

Ex- vivo tissues from control mice and mice treated with doxycycline to induce overexpression of VAV1-L1CAM-CXCR4 were analyzed immediately after dissection using the IVIS Spectrum imaging system (PerkinElmer). Prior to sacrifice, mice were injected intraperitoneally with luciferin (150 mg/kg body weight) to enable bioluminescence detection. Following luciferin injection, mice were euthanized, and the organs of interest (liver, spleen, lungs, and kidneys) were harvested. Bioluminescence imaging was performed using the IVIS Spectrum imaging system, and the resulting images were analyzed to quantify the bioluminescent signals. For each organ, five regions of interest (ROIs) were randomly defined within the tissue samples. Background luminescence was subtracted from each ROI measurement to obtain normalized luminescence intensities. Bioluminescence data were then visualized using violin plots with datapoints representing multiple ROI measurements per organ for both control and Dox-treated mice across the four different organs. Statistical analyses were conducted using PRISM software (GraphPad Software).

### Microscope imaging

mCherry fluorescence and brightfield images were acquired using a Leica DM Fluo inverted fluorescence microscope equipped with appropriate filter sets for mCherry. Prior to imaging, samples were prepared according to standard protocols. Fluorescence images were captured using the appropriate excitation and emission filters, while brightfield images were acquired simultaneously to provide structural context.

### Western blot

Western blotting analysis was performed using standard methods. Whole-cell extracts were prepared in lysis buffer [50 mM tris (pH 8.0), 50 mM NaCl, 1.0% NP-40, 0.5% sodium deoxycholate, and 0.1% SDS] containing protease inhibitor cocktail (cOmplete, Roche) and phosphatase inhibitor cocktail (Thermo Fisher Scientific). Equal amounts of protein, as determined by the Micro BCA Protein Assay Kit (Pierce), were resolved on NuPage Novex 4 to 12% bis-tris gels (Invitrogen) or NuPAGE Novex 7% tris-acetate protein depending on the protein size and transferred onto nitrocellulose membranes (0.2 μm, Whatman).

Membranes were blocked in phosphate-buffered saline with 0.1% Tween 20 (PBST) 5% bovine serum albumin (BSA) for 1 hour, incubated with primary antibodies in PBST 1% BSA overnight at 4°C and with secondary antibodies coupled to horseradish peroxidase for 45 min in PBST 1% BSA. Bands were visualized using an enhanced chemiluminescence detection reagent (GE Healthcare). Primary antibodies used against the following antigens were as follows: anti-vimentin D21H3 rabbit monoclonal antibody (mAb); Cell Signaling Technology, #5741], anti–E-cadherin (24E10 rabbit mAb; Cell Signaling Technology, #3195), anti–N-cadherin (rabbit mAb; Cell Signaling Technology, #4061), anti–Sox2 (rabbit polycolonal Abcam, #15830),anti-slug(rabbit mAb; Cell Signaling Technology, #4933) anti-snail (rabbit mAb; Cell Signaling Technology, #4933) anti-ZO1 (rabbit mAb; Cell Signaling Technology, D7D12 #8193) anti-RAS(rabbit mAb; Cell Signaling Technology, #67648)anti-slug(rabbit mAb; Cell Signaling Technology, #4933) anti-vinculin (mouse clone h-VIN1; Sigma-Aldrich, #V9131), anti–FOXA1 (Abcam AB23738 rabbit mAb; anti-FOXP1 (rabbit mAb; Cell Signaling Technology, #2005), anti–glyceraldehyde-3-phosphate dehydrogenase (GAPDH) (rabbit mAb; Santa Cruz Biotechnology, #sc-25778), anti-H3K273me [rabbit polyclonal antibody (pAb); Cell Signaling Technology, #9733], anti-total H3 (rabbit mAb; Cell Signaling Technology, #AB1791), anti-CAS9 polyclonal Diagenode #C15310258), anti–c-JUN (60A8 rabbit mAb; Cell Signaling Technology, #9165), anti–Ras (G12D Mutant) Recombinant Rabbit Monoclonal Antibody (HL10)Thermofisher MA5-36256, anti tubulin monoclonal Sigma #T5168

### Library cloning and amplification

IFN library cloning was performed in the CROP-SEQ-mCherry2-Puro as described in Datlinger et al, 2017. Briefly, the modified version of CROPseq-Guide-Puro containing the fluorescent protein m-Cherry was digested BsmBI. The Assembly of gRNA-encoding ssDNA oligos into the vector backbone by Gibson Assembly. The library-plasmid amplification was performed by electroporation of Lucigen Endura cells (64242-2-LU). Next generation sequencing of gRNA sequences for library quality control or pooled screens.

Single sgRNA targeting KRAS and all the genes used for validation control have been cloned using the same strategy to clone the library. All the sgRNAs sequences are indicated in supplementary figure 3.

The original Cropseq Cherry Puro vector was modified to replace the cherry fluorescent marker with blue fluorescent protein (BFP). This vector was then used to clone single guide RNAs (sgRNAs) targeting KRAS, enabling the interference of KRAS expression levels.

### Lentivirus production and infection

Lentivirus was produced by transfecting human embryonic kidney–293 T cells using FuGENE HD (Promega) as described (Gargiulo 2013). Supernatant infection of IFN library in KPEluc-dcas9vpr cells was performed as described in the references above, and NSCLC cells were infected separately by one round of overnight exposure to the viral pool. Multiplicity of infection was experimentally designated as <0.5 based on mCherry2 expression upon low-MOI infection.

### In vivo CRISPRa screening procedure

A total of 1 × 10˄6 KPE-lucTETON-dCas9-VPR-IFN library-mcherry were injected intravenously via the tail vein (500x representation). Successful engraftment was confirmed using noninvasive bioluminescence imaging. Tumor growth in the lungs was monitored using an IVIS imaging system. IVIS Lumina imaging was performed as described previously (Gargiulo et al., 2014). After 7 days post-injection, mice were randomized and were treated or not with doxycycline administered in the drinking water. Tumor growth in the lungs and distant organs was monitored regularly using IVIS imaging. Animals showing signs of respiratory distress were euthanized, and primary lung tumors, liver, kidney, bones, and brain tissues were dissected. Primary cells were isolated from these tissues. CTCs were isolated from the retroorbital vein. Isolated cancer cells were analyzed for mCherry expression using fluorescence-activated cell sorting (FACS).

### CRISPR activation and fate mapping procedure

Total RNA was extracted using the Kit Stratec. The concentration of the RNA was quantified by the Qubit RNA HS Assay Kit (Invitrogen). The integrity of the RNA was determined with the High-Sensitivity RNA ScreenTape System (Agilent). 45 nanograms of total RNA per sample was used as input for constructing CROP-seq transcript-specific multiplexed 3′-cDNA (complementary DNA) libraries using modified barcoded primers ^62^ in an adapted version of bulk RNA barcoding and sequencing protocol ^63^. The final multiplexed library pools were quantified with the Qubit dsDNA HS Assay Kit (Invitrogen) and the Collibri Library Quantification Kit (Invitrogen), and the proper PCR library fragment size was assessed by the TapeStation High-Sensitivity D1000 ScreenTapes Kit (Agilent). Sequencing of the pooled libraries was performed on NovaSeq 6000 in a paired-end mode (read 1, 21 bp; index i7, 8 bp; index i5, 8 bp; read 2: 150 bp). The initial demultiplexing based on Illumina indices was performed using the bcl2fastq conversion software (v2.20.0). Next, the CROP-seq barcode sequences were extracted from read 1 reads using cutadapt (v2.1) and used for subsequent internal demultiplexing using BRBseqTools-1.6.jar ( http://github.com/DeplanckeLab/BRB-seqTools). The 20 bp sgRNA sequences were extracted from demultiplexed reads using cutadapt (v2.1) and aligned to the reference custom IFN library using bowtie2 (v2.4.4), for subsequent generation of sgRNA read counts. The initial quality control and exploratory analysis were performed in the R 4.1.2 environment. First, screen samples with shallow library coverage (<100,000 recovered reads) were omitted from the analysis. The sgRNAs detection rate was calculated for each sample, providing the basis for defining the clonality of the metastatic samples, with the detection rates between the 0.25 and 0.5 quantiles defining the oligoclonal, and those above the 75th quantile – the polyclonal metastatic samples. The read counts were further median scaled (via normalizeMedianValues function, gCrisprTools package (v.2.0.0) and used to define stable groups for differential analysis. As a comparison measure sample diversity was estimated by calculating Shannon entropy index of the respective sample sgRNAs frequencies. Further, for each group of samples (controls, primary tumor, and poly-/oligo or tissue-specific metastasis) inter-group correlation (Pearson correlation, average linkage) was calculated. Samples with the lowest mean correlation scores were iteratively left out when necessary until cluster stability was achieved. The group core samples were subjected to MAGeCK (v.0.5.9.3) test command, with “--norm-method median --control-sgrna --remove-zero control --remove-zero-threshold 50 --sort-criteria pos” parameters specified. For the metastatic samples with median sgRNA abundance of zero, a pseudocount was added to the sgRNA count for each sample included in the comparison. Enriched genes at the FDR threshold of 0.05 were considered significant.

### In vivo competition assay

For in vivo validation, we conducted a competition assay by injecting a mixture of KPE-luc dCas9-VPR containing 7 sgRNAs cloned in the CRISPR-Cas9 knockout pooled library (CropSeq) tagged with mCherry (40%) and KPE-dCas9-VPR-luc GW human library (60%) via the tail vein (total cells injected 2×105). Tumor growth was monitored using an IVIS imaging system. Subsequently, doxycycline was administered in the drinking water, and primary tumors and metastatic tissues were harvested at the humane endpoint. Primary cells isolated from these tissues were analyzed using flow cytometry (FACS) to assess the expression levels of BFP and mCherry.

### Genome editing of cancer cells

Genome editing of FOXP1 and FOXA1 of all cell lines described was performed by electroporation using the Amaxa 4D-Nucleofector Kits and Kit v2 by Synthego. Briefly, 2 × 10^5^ cells were counted and resuspended in 20 μl of the respective buffers and supplement in a 16-well Nucleocuvette strip. For KPE electroporation was performed with P3 nucleofection buffer (Lonza) and using CM150. Editing efficiency was estimated by western blot and Sanger sequencing and calculated using the ICE (inference of CRISPR edits) webtool provided by Synthego.

### ATAC sequencing

ATAC-seq was performed on primary lung and metastatic isolated KPE cells treated with or without IFN (+ 10ng/ml TNFa + 5ng/ml IFNg + 5ng/ml IL1B + 2ng/ml OSM (1µl of) medium and TGF-β1 (5 ng/ml). A total of 60,000 cells were sorted in biological replica and centrifuged; the pellet was gently resuspended in 50 μl of ATAC mix [25-μl 2× tagmentation DNA (TD) buffer, 2.5-μl 891 transposase and 22.5-μl nuclease-free water from Nextera DNA Library Prep, Illumina]. Cells were incubated for 60 min at 37°C with moderate shaking (500 to 800 rpm), lysed in proteinase K and AL buffer (QIAGEN); DNA was purified using 1.8× AMPure XP beads (Beckman Coulter). Library prep was made using primers compatible with Nextera Illumina. Each library was individually quantified using Qubit 3.0 Fluorometer (Life Technologies) and profiled on a TapeStation High Sensitivity D1000 ScreenTape (Agilent). The multiplexed libraries were sequenced on a HiSeq 4000 in a 2× 75–base pair (bp) mode.

### RT–qPCR and RNA-seq

RNA was isolated using Trizol and cDNA was generated using SuperScript II or VILO according to the manufacturer’s instructions (Invitrogen). Primer details are available upon request. For RNA-seq, the library was prepared using Brb seq RNA sample protocol.

### RNA seq

RNA was extracted using the STATEC kit. The concentration of the RNA was quantified by the Qubit RNA HS Assay Kit (Invitrogen). The integrity of the RNA was determined with the High-Sensitivity RNA ScreenTape System (Agilent). Sixty nanograms of total RNA per sample was used as input for constructing multiplexed 3′-cDNA (complementary DNA) libraries using barcoded oligo-dT primers (*<u>44</u>*) in an adapted version of bulk RNA barcoding and sequencing protocol (*<u>45</u>*). The final multiplexed library pools were quantified with the Qubit dsDNA HS Assay Kit (Invitrogen) and the Collibri Library Quantification Kit (Invitrogen), and the proper library fragment distribution was assessed by the TapeStation High-Sensitivity D1000 ScreenTapes Kit (Agilent). Sequencing of the pooled libraries was performed on NovaSeq 6000 in a paired-end mode (read 1, 21 bp; index i7, 8 bp; index i5, 8 bp; read 2: 150 bp). The initial demultiplexing based on Illumina indices was performed using the bcl2fastq conversion software (v2.20.0). Next, the oligo-dT barcode sequences were extracted from read 1 reads using cutadapt (v2.1) and used for subsequent internal demultiplexing using BRBseqTools-1.6.jar (http://github.com/DeplanckeLab/BRB-seqTools). The demultiplexed data were aligned to a mouse GRCm38.102 genome using STAR (v2.6.0c), and the count matrices were subsequently generated using HTSeq (v2.0.2).

### Pro-metastatic upstream regulators cytokine screening

KPEluc-dCas9VPR were seeded in basal medium or in IFN medium containing 10ng/ml TNFa, 5ng/ml IFNg 5ng/ml IL1B + 2ng/ml OSM).

Cells were treated with cytokines for 24 hours at two varying concentrations with 10-fold difference (low and high concentration). Post-treatment, cells were fixed with 1% PFA for 10min, washed with PBS and stained for NCAM1 to assess staining patterns through a high-content imaging cytokine screen. Permeabilization and blocking were performed using a solution containing 4% BSA and 0.1% Triton-X in 1x PBS with 30 min incubation at room temperature, followed by one wash with PBS. Primary antibody staining was performed using anti-NCAM1 (mouse, Sigma, T5168) at a 1:2000 dilution and pre-conjugated Phalloidin-FITC. The primary antibodies were diluted in a solution of 4% BSA-PBS with 0.1% Triton-X and added to the wells, excluding control wells that served as secondary-only controls. The plates were incubated for 1 hour at room temperature in the dark, followed by two washes with PBS. Secondary antibody staining was carried out using Alexa 647 (anti-rabbit, Invitrogen, A31573), diluted at 1:200 in 0.5% BSA in 1x PBS. The secondary antibody was incubated for 45 min at 4°C in the dark, followed by two washes with PBS. For nuclear staining, Hoechst was prepared at a 1:1000 dilution in 1x PBS and incubated for 5 minutes, following three washes with PBS. Imaging was conducted using the Operetta high-content imaging system (Revvity) in confocal mode with a 40x water objective, capturing 25 fields per well. Using the secondary only conditions, appropriate filter and exposure settings were determined to image Alexa647, FITC, Hoechst33258 and brightfield channels. A z-stack of three planes with 2µm increments was captured. Images were analyzed by creating maximum projections of the z-stacks. Each channel was filtered using a rolling ball algorithm with a radius of 10 pixels. Nuclei were identified using the inbuilt image detection algorithms (Method A) on the Hoechst-stained images, and cells were identified using Method F on Alexa647 (Tubulin) and FITC (Actin-Phalloidin-stained) images. Morphology properties were calculated using the STAR Methods, recording mean and standard deviation parameters of cell area, roundness, width, length, compactness threshold and radiality. All derived cell morphology parameters were used to segment single cells and read out NCAM1 staining intensities per cell. Data is presented as bubble plots, with the number of nuclei per condition representing bubble size and the fold-change induction of NCAM1 staining per concentration on the x-axis. Statistical analysis used multiple comparisons of NCAM1 intensity means with Dunnett contrasts.

### Fluorescence-activated cell sorting (FACS)

Harvested single cell suspensions were resuspended in cold media and filtered into FACS tubes. FACS analysis was performed on a BD LSR Fortessa system. Sorting was carried out on a BD ARIA III system. Depending on the fluorophores to be analysed, the suitable laser-filter combinations were determined and acquired. To exclude dead cells, events were typically gated according to their shape and granularity (FSC-A vs. SSC-A) and doublets were removed (FSC-A vs. FSC-H). Positive gates were set above background thresholds with low to zero levels of the fluorophore of interest using unstained or wildtype cells. FlowJo v10 was used for all analyses.

### Single cell RNA-seq analysis

The previously published patient advanced non-small cell lung cancer and metastatic lung adenocarcinoma single-cell RNA-seq datasets were retrieved from the Gene Expression Omnibus database via respective accession codes (GSE148071, GSE123904) and analyzed separately using the Seurat toolkit (v4.3.0) in R (4.1.2). Standardly, the obtained gene expression matrices were filtered removing cells with less than 200 and more than 5000 expressed genes, as well as cells with a mitochondrial content higher than 30%. doubletFinder (v2.0.3) was used to discriminate doublets and copykat (v1.0.5) was used to infer genomic copy numbers from the respective single-cell RNA-seq datasets. Individual patients single-cell RNA-seq profiles were integrated using the FindIntegrationAnchors() and IntegrateData() functions (normalization.method = “SCT”). PCA and UMAP dimensionality reduction were performed on normalized scaled datasets, cluster identification was performed using FindClusters() function and cell type annotation was conducted using the ScType package.

### Human Bulk RNA-seq Analysis

Data Integration. Processed RNA-seq, somatic mutation, and clinical data from TCGA were downloaded via TCGAbiolinks. KRAS variants classified as “Missense_Mutation” were considered KRAS mutants. Processed RNA-seq, somatic mutation, and clinical data from PRISM were downloaded from Gustave Roussy’s cBioPortal ([https://cbioportal.gustaveroussy.fr/study/summary?id=metaprism_2023](https://cbioportal.gustaveroussy.fr/study/summary?id=metaprism_2023)) and nextcloud server ([https://nextcloud.gustaveroussy.fr/s/HFB6QgycJ56EpiC](https://nextcloud.gustaveroussy.fr/s/HFB6QgycJ56EpiC)). KRAS variants classified as “Missense_Mutation” were considered KRAS mutants. Kallisto-generated raw RNA-seq counts were rounded to obtain integer raw counts for downstream analysis. HMF (Hartwig Medical Foundation) RNA-seq, somatic mutation, and clinical data were accessed via [Hartwig Medical Foundation](https://www.hartwigmedicalfoundation.nl/en/) data access request procedure. Processed data were used in the analysis. KRAS variants classified as “Missense_Mutation” and putative driver were considered KRAS mutants. Processed data from TRACERx were downloaded from [https://zenodo.org/records/7822002](https://zenodo.org/records/7822002), [https://zenodo.org/records/7649257](https://zenodo.org/records/7649257), and [https://zenodo.org/records/7603386](https://zenodo.org/records/7603386). KRAS variants classified as “nonsynonymous SNV” and putative driver were considered KRAS mutants. RSEM-generated raw RNA-seq counts were rounded to obtain integer raw counts for downstream analysis. LUAD, LUSC, PAAD, and COREAD (COAD + READ) samples were used from TCGA, PRISM, and HMF, while LUAD and LUSC samples were used from TRACERx.

Gene IDs of raw RNA-seq count data from all datasets were mapped, retaining 18,727 protein-coding genes available in all datasets, and datasets were merged (supplementary material).

Since lung cancer samples from HMF were originally classified as only NSCLC (i.e., not sub-classified as LUAD or LUSC), HMF-NSCLC samples were classified into LUAD or LUSC using PRISM and TRACERx LUAD and LUSC data. From differential expression results between LUAD and LUSC (with DESeq2 [Love et al.] v1.42.1, ‘design = ∼ source + diseasè, 13,674 genes, excluding 5,053 genes in downstream 95 gene sets and KRAS out of 18,727 total integrated protein-coding genes to exclude possible variable genes, were used to calculate size factors with the ‘DESeq2::estimateSizeFactors’ function), upregulated top 150 genes for each LUAD and LUSC were identified (LUAD_genes, LUSC_genes). Then single-sample enrichments of LUAD_genes (LUAD_enrichment) and LUSC_genes (LUSC_enrichment) were calculated via the GSVA method [Hänzelmann et al.] (GSVA v1.50.5 ‘kcdf = “Gaussian”’) with VST (variance stabilizing transformation) values of datasets calculated separately via the ‘DESeq2::varianceStabilizingTransformation’ function (with ‘blind = FALSE, design = ∼ disease_KRAS_mutation_status’). Then a first-degree support vector machine classifier was fit using LUAD_enrichment and LUSC_enrichment as input to classify HMF-NSCLC samples into LUAD or LUSC (supplementary material). After data merging, batch effect correction between sources (TCGA, PRISM, HMF, TRACERx) was performed with ComBat-seq [Zhang et al.] (’batch = source, covar_mod = model.matrix(∼ stage + disease + kras_mutation_status)‘) by exploiting putative similarities between biological groups (i.e., the same cancer type in the same stage should be similar across sources, see supplementary material). After batch correction, VST was performed with the following settings using DESeq2: Top 150 stable genes from Bhuva et al. were used to calculate size factors with the ‘DESeq2::estimateSizeFactors’ function, and then VST was performed with the ‘DESeq2::varianceStabilizingTransformation’ function (with ‘blind = FALSE, design = ∼ stage + disease + KRAS_mutation_status’). The VST expression values were used in Cox proportional hazards models and KRAS expression level assignment. To define KRAS expression level of a sample (either KRAS-high or KRAS-low), the following approach was implemented: Patients were divided into groups based on the combination of data source (TCGA, PRISM, HMF, or TRACERx), disease (LUAD, LUSC, PAAD, or COREAD), and KRAS mutation status (mutant or wild type). Then median KRAS VST expression value was calculated for each group, and samples were classified into KRAS-high or KRAS-low with respect to their group’s median KRAS expression. The KRAS-level of a sample and grouping of samples defined in this step were used in differential expression analysis.

All human bulk RNA-seq data integration and analysis was performed in R 4.3.3 environment.

### Survival Analysis

All Cox proportional hazards models and Kaplan-Meier curves were generated in R 4.3.3 using the survival and survminer packages. The log-rank test was used to determine the statistical significance of Kaplan-Meier curves.

In KRAS-level Kaplan-Meier analysis, KRAS-level (KRAS-high or KRAS-low) was defined in the following way: Patients were divided into groups based on the combination of data source (TCGA, PRISM, HMF) and KRAS mutation status (mutant or wild type). Then median KRAS log2(TPM+1) expression value was calculated for each group, and samples were classified into KRAS-high or KRAS-low with respect to their group’s median KRAS expression. TPM (transcript per million) values come from data source, no batch correction was applied to original TPM values since Kaplan-Meier analysis was performed on within-dataset level.

RAS84 Index (RI) values were calculated as the mean VST value of RAS84 genes [East et al.]. Here VST expression values were calculated on within-dataset level in the following way: 13,674 genes (excluding 5,053 genes in downstream 95 gene sets and KRAS out of 18,727 total integrated protein-coding genes to exclude possible variable genes) were used to calculate size factors with the ‘DESeq2::estimateSizeFactors’ function. Patients were divided into groups based on the combination of disease (LUAD, LUSC, PAAD, or COREAD), and KRAS mutation status in every dataset separately (TCGA, PRISM, HMF). And then ‘DESeq2::varianceStabilizingTransformation’ function were used for VST calculation with ‘blind = FALSE, design = ∼ group’. Then median RI value was calculated for each group, and samples were classified into RI-high or RI-low with respect to their group’s median RI.

### Gene Set Enrichment Analysis

All gene set enrichment analyses were performed with the ‘fgsea’ R/Bioconductor package v1.28.0 [Korotkevich et al.] using Wald statistics from DESeq2 differential expression results of desired contrasts (i.e., comparisons between selected groups).

MSigDB Hallmark Gene Sets were retrieved via msigdb R/Bioconductor package. ‘Cilia Signature’ was derived from Patir et al., filtering their signature by: Human Protein Atlas Bronchus Cilia staining as “Positive Stain” and FOXJ1 gene expression profile as “Associated”. ‘AT1 Signature’ and ‘AT2 Signature’ were derived from AT1 and AT2 signature lists of Li et al., Travaglini et al. and Sikkema et al. All genes and gene sets used in human bulk RNA-seq analysis are provided in supplementary material 1.

### Differential Expression Analysis

ComBat-seq batch-corrected integer expression values (see above) were used for differential expression analysis with DESeq2 with ‘design = ∼ stage * kras_status * kras_level’. 150 stable genes (see above) were used for the ‘DESeq2::estimateSizeFactors’ function.

Differential expression analysis of longitudinal TRACERx LUAD KRAS mutant samples was performed in the similar fashion but with paired design ‘design = ∼ patient + kras_status * kras_level * stagè.

In figure S7B where different metastatic tissues (lymp node, metastatic lung or distant organ) were compared to primary lesion side, ‘tissue_typè variable were used in the designs instead of stage, such that ‘design = ∼ tissue_type * kras_status * kras_level‘, and for longitudinal analysis of KRAS mutant TRACERx data ‘design = ∼ patient + kras_level * tissue_typè.

### PROGENy Pathway Activity Inference

PROGENy pathway activity inference was performed using the decoupleR R/Bioconductor package v2.8.0 [Badia-I-Mompel et al.]. The PROGENy [Schubert et al.] model was retrieved with ‘decoupleR::get_progeny(organism = ‘human’, top = 1500)’ from OmnipathR v3.15.1. Model was fit with the ‘run_mlm’ function in contrast-based mode using Wald statistics from DESeq2 differential expression results of indicated contrasts.

### DepMap Data Analysis

DepMap[Tsherniak et al.] data (Version: DepMap Public 24Q2) were obtained from [DepMap’s data portal](https://depmap.org/portal/data_page/?tab=allData). For evaluating KRAS-high and KRAS-low signature enrichment across CCLE/DepMap cell lines, we selected 316 cell lines representing 9 cancer types: LUAD, PAAD, COAD, LUSC, STAD, ESCA, IHCH, PLBMESO, and BLCA. To score these cell lines, we used the file “OmicsExpressionProteinCodingGenesTPMLogp1BatchCorrected.csv” from DepMap.

Differential expression analysis results for “Mutant, KRAS-high vs. KRAS-low” and “Metastatic Mutant, KRAS-high vs. KRAS-low” were integrated in a majority-vote framework to create six gene sets (signatures): three for KRAS-high and three for KRAS-low, with gene thresholds set at 100, 300, and 500 to minimize the impact of gene set size.

For each signature, a majority vote was based on ranking the sum of integrated statistics (-log10(padj) * log2(FoldChange)). We selected the top-ranking genes for KRAS-high and the lowest-ranking genes for KRAS-low to form each gene set. Gene Set Variation Analysis (GSVA) was applied to score the cell line expression profiles, and the mean values of the 100, 300, and 500 gene set enrichments were calculated to derive the final KRAS-high or KRAS-low enrichment scores (Fig. S7C).

Finally, the difference between KRAS-high and KRAS-low scores was computed for each cell line, and these differences were scaled. The resulting scores are shown on the y-axis in Fig. 7G.

### MetMap Data Analysis

MetMap500[Jin et al.] data were accessed from [DepMap’s MetMap data portal](https://depmap.org/metmap/data/index.html) and integrated with DepMap data to incorporate information on KRAS expression, mutation status, and copy number alterations. Pearson correlations between KRAS expression and mean metastatic potential were calculated across various subsets based on KRAS mutation status and copy number alterations, as shown in Fig. 3G.

### KP-Tracer Data Analysis

Processed KP-Tracer scRNA-seq data were obtained from [Zenodo](https://zenodo.org/records/5847462) [Yang et al.]. In Figs. 3F and S3H, we classified cell states and fates according to the authors’ framework, defining cell states in “Fate Cluster 3” as “late” and those in “Fate Clusters 1 and 2” as “early.”

For contrast-based PROGENy (Fig. S5D) and CollecTRI (Fig. 6H) analyses, we first created pseudobulk profiles by aggregating raw counts through summation. We then conducted differential expression analysis using DESeq2 with a ‘design = ∼ stagè model for the “Late vs. Early” comparison, and ‘design = ∼ stage_level’ for other comparisons.

Pathway activity inference was carried out using the PROGENy model through the decoupleR R/Bioconductor package (v2.8.0) [Badia-I-Mompel et al.]. We retrieved the PROGENy model with ‘decoupleR::get_progeny(organism = ‘mouse’, top = 500)‘ from OmnipathR (v3.15.1) and fitted the model in contrast-based mode with the ‘run_mlm’ function, utilizing Wald statistics from DESeq2 differential expression results for the specified contrasts.

Similarly, transcription factor activity inference was performed with the CollecTRI model [Müller-Dott et al.], also using decoupleR (v2.8.0). The CollecTRI model was retrieved via ‘decoupleR::get_collectri(organism = ‘mouse’, split_complexes = FALSE)‘ from OmnipathR (v3.15.1) and fitted in contrast-based mode with the ‘run_ulm’ function, applying Wald statistics from DESeq2 differential expression results for each contrast.

### Gene Set Enrichment Analysis of cell line-derived xenografts

Bulk RNA-Seq from PDX models engrafted with lung cancer cell lines treated with KRAS-G12C inhibitor MRTX849 were downloaded from SRA database (project number PRJNA578935). Low-quality reads and adaptors were removed by fastp v.0.23.2 (parameters --average_qual 30 --length_required 100 --detect_adapter_for_pe) and xengsort v.2.0.5 was used to remove mouse reads. Good quality reads were pseudoaligned against the human transcriptome (v.30 Ensembl 96) by kallisto v.0.44.0 (parameters-b 100). Counts were loaded into R v.4.3.2 and lowly expressed genes removed prior to differential expression analysis (minimum of 10 reads in at least one sample and a minimum of 20 reads across all samples). Data was normalized and differentially expressed genes were identified using DESeq2 v.1.42.1. The function GSEA from package clusterProfiler v.4.10.0 and the Molecular Signature Database v.2023.2 were used for Gene Set Enrichment Analysis on hallmark gene sets and the cilia signature. KRAS copy number was from https://depmap.org/portal (DepMap 24Q2 Public. Figshare+. Dataset. https://doi.org/10.25452/figshare.plus.25880521.v1) and allele frequency from Fastp - https://doi.org/10.1093/bioinformatics/bty560 Xengsort - https://doi.org/10.1186/s13015-021-00181-w Kallisto - https://doi.org/10.1038/nbt.3519 DESeq2 - https://doi.org/10.1186/s13059-014-0550-8 clusterProfiler - https://doi.org/10.1016/j.xinn.2021.100141 MsigDB - https://doi.org/10.1073/pnas.0506580102

### Patient-derived xenografts

The lung cancer xenograft model was obtained from the Victorian Cancer Biobank. The patient-derived xenograft (PDX) model was generated from a patient with a lung adenocarcinoma that contained the following mutation in KRAS (c.34G>T; p.G12C). Ethical approval was obtained to generate this model. In collaboration with TRACE the PDX model was further expanded in Leuven and F4 generations were used for all experiments.

### KRAS inhibition in vivo

When the tumors reach a volume of 300 mm^3^, the PDX mice were injected daily with 100mg/kg of the KRAS inhibitor MRTX849 dissolved in citrate buffer (0,05M pH5) and with 10% captisol via oral gavage. Control mice were injected with 0,05M citrate buffer only. After 67 days of treatment the drug administration as stopped, and the tumor size was measured regularly. Random mice were sacrificed before treatment and after 7 or 20 days of treatment and the tumors were embedded for further analysis.

### OPAL staining

An OPAL-based approach, which relies on individual tyramide signal amplification (TSA)- conjugated fluorophores to detect various targets, was used to perform the staining. Sections (5 mm) of formalin-fixed, paraffin-embedded tumors were deparaffinized and subjected to antigen retrieval in citrate buffer pH6. Blocking was performed for 30 minutes with 10% goat serum in 1% BSA and 01% Triton X-100 in PBS. The sections were incubated overnight at 4°C with the following primary antibodies anti-acetylated tubulin (Merck, T7451, 1/2000); anti-Arl13b (Sanbio, 17711-1-AP, 1/500) and anti-FANK1 (Merck, HPA038413, 1/50). After washing with TBST, the sections were incubated for 15 min at room temperature with the appropriate HRP and Fluor tyramides (PerkinElmer) to detect antibody staining, prepared according to the manufacturer’s instructions: Opal 570 or OPAL 690 (dilution 1:150). Stripping of primary and secondary antibodies was performed by placing the slides into a plastic container filled with antigen retrieval (AR) buffer in citrate buffer pH 6. A microwave was used to heat the liquid to 100 °C (2 min), and the sections were then microwaved for an additional 15 min at 75 °C. Slides were allowed to cool down in the AR buffer for 15 min at room temperature and were then rinsed with deionized water and 13 Tris-buffered saline with Tween-20. After three additional washes in deionized water, the slides were counterstained with DAPI for 5 min and mounted with ProLong Gold Antifade Mountant (Thermo Fisher Scientific, P36930).

### Statistics & Reproducibility

Unless otherwise indicated, the standards of the analyses were as follows. For multiple comparisons of two groups, unpaired two-tailed Student’s t-test was used, unless otherwise specified. For comparisons of two or more groups, one-way ANOVA followed by Dunnett’s post-hoc multiple comparisons test correction. For correlation analyses, Pearson correlation coefficients were calculated. Hierarchical clustering used Manhattan distance calculations. For boxplot representations, data distribution is shown with box indicating the interquartile range and inner line indicating the median. Whiskers extend to represent the data range, including outliers. Barplot data is shown as mean value +/- standard deviation. All experimental data has been derived from at least three independent biological replica. No data that passed QC were excluded from the analyses and the investigators were not blinded to allocation during experiments and outcome assessment.

## Acknowledgments

We are grateful to Christoph Bock (CeMM) for sharing the CROP-seq-mCherry prior to publication. We are also grateful for excellent assistance to Carlos Company, Giorgia Di Benedetto, Jonas Eberle, Melanie Grossmann, Marialucia Massaro and Jikke Wierlix (MDC, technical assistance), to T. Borodina, T. Conrad, J. Wilde, Madlen Sohn, Daniele Sunage-Franze, J. Altmuller from the MDC sequencing platforms (sequencing experiments), to S. Dietrich (MDC) and M. Richter and A. Spotbert from the MDC ALM platform ( lightsheet microscopy), Greet Bervoets (VIB, PDX IHC staining), Yoann Pradat and Sergey Nikolaev (Gustave Roussy, META-PRISM) Nicholas McGranahan, Piotr Pawlik and Tom Jones (UCL, TRACERx data). Data analyses include data generated by the TCGA Research Network: https://www.cancer.gov/tcga, and by the HMF (accessed under a license agreement DR-332). The data and lung cancer tissue used in the creation of the Xenografts was provided by the Victorian Cancer Biobank (AUS) with appropriate ethics approval. The Victorian Cancer Biobank is supported by the Victorian Government. This project was supported by DFG (DFG SE 2847/2-1 to M.S.). The G.G. lab acknowledges funding from MDC, Helmholtz (VH-NG-1153), ERC (714922). The PDX work was funded by Boehringer Ingelheim RCV & Co KG to J-C. M. Y.D. is a graduate students with Humboldt University and S.K. with Charitè Medical University. S.K. and Y.D. are affiliated with the Berlin School of Integrative Oncology (BSIO) at Charitè.

## Deposited data and code

The whole-genome sequencing, RNA sequencing and corresponding clinical data used in this study were made available by the TCGA (phs000178.v3.p3, http://cancergenome.nih.gov), by META-PRISM (https://cbioportal.gustaveroussy.fr:8081/study/summary?id=metaprism_2023), and TRACERx (https://zenodo.org/records/7822002, https://zenodo.org/records/7649257, https://zenodo.org/records/7603386). All the relevant data generated here were deposited in the appropriate repositories. CRISPRa, RNA-seq and ATAC-seq data were uploaded at GEO (GSE272096).

A secure token has been created to allow review of record GSE272096 while it remains in private status: ujejwkeifzczrsj For additional requests about data availability please contact the corresponding authors.

## Code availability

No new custom code or mathematical algorithm that is deemed central to the conclusions was developed in this study. All data were processed and analyzed building on existing codes and softwares as detailed in the Methods section. For additional requests about codes availability please contact the corresponding authors.

## Supplementary data for peer review

Aggregated counts for CRISPRa screen (Fig.1 and Fig.2 key data) are provided to Reviewers in Supplementary Table S1_CRISPRa-screen-source-data.txt. Cox Proportional Hazards in TCGA cohort are provided in Supplementary_Table_S2_coxph_primary_luad_samples.xlsx (Fig. 4 key data). Fig. 7 key data are provided as Supplementary_Table_S3_Human-bulk-RNA-seq-source-data.xlsx (GSEA) and Supplementary_Table_S4_LUAD_vs_LUSC_classification.xlsx (NSCLC sub-classification).

## OPAL staining

An OPAL-based approach, which relies on individual tyramide signal amplification (TSA)- conjugated fluorophores to detect various targets, was used to perform the staining. Sections (5 μm) of formalin-fixed, paraffin-embedded tumors were deparaffinized and subjected to antigen retrieval in citrate buffer pH6. Blocking was performed for 30 minutes with 10% goat serum in 1% BSA and 01% Triton X-100 in PBS. The sections were incubated overnight at 4°C with the following primary antibodies anti-acetylated tubulin (Merck, T7451, 1/2000); anti-Arl13b (Sanbio, 17711-1-AP, 1/500) and anti-FANK1 (Merck, HPA038413, 1/50). After washing with TBST, the sections were incubated for 15 min at room temperature with the appropriate HRP and Fluor tyramides (PerkinElmer) to detect antibody staining, prepared according to the manufacturer’s instructions: Opal 570 or OPAL 690 (dilution 1:150). Stripping of primary and secondary antibodies was performed by placing the slides into a plastic container filled with antigen retrieval (AR) buffer in citrate buffer pH 6. A microwave was used to heat the liquid to 100 °C (2 min), and the sections were then microwaved for an additional 15 min at 75 °C. Slides were allowed to cool down in the AR buffer for 15 min at room temperature and were then rinsed with deionized water and 13 Tris-buffered saline with Tween-20. After three additional washes in deionized water, the slides were counterstained with DAPI for 5 min and mounted with ProLong Gold Antifade Mountant (Thermo Fisher Scientific, P36930).

## Authors’ Contributions

M. Serresi: Conceptualization, resources, data curation, formal analysis, supervision, funding acquisition, validation, investigation, visualization, methodology, project administration, writing-review and editing. S. Kertalli: Validation, investigation, visualization, and methodology. Y. Dramaretska: data curation, formal analysis, validation, investigation, visualization, methodology. M. J. Schmitt: formal analysis, validation, visualization, methodology. A. O. Cetin: data curation, formal analysis, validation, investigation, visualization, methodology. H. Naumann and M. Zschummel: methodology. M. Liesse-Labat: resources. L. F. Maciel: validation, visualization, methodology. J. Declercq: validation, investigation, visualization, methodology. M. Samwer: resources. J-C. Marine: resources, data curation, formal analysis, supervision, funding acquisition, writing–review and editing.

G. Gargiulo: Conceptualization, resources, data curation, formal analysis, supervision, funding acquisition, validation, investigation, visualization, methodology, writing-original draft, project administration, writing–review and editing.

---

**Figure S1.**
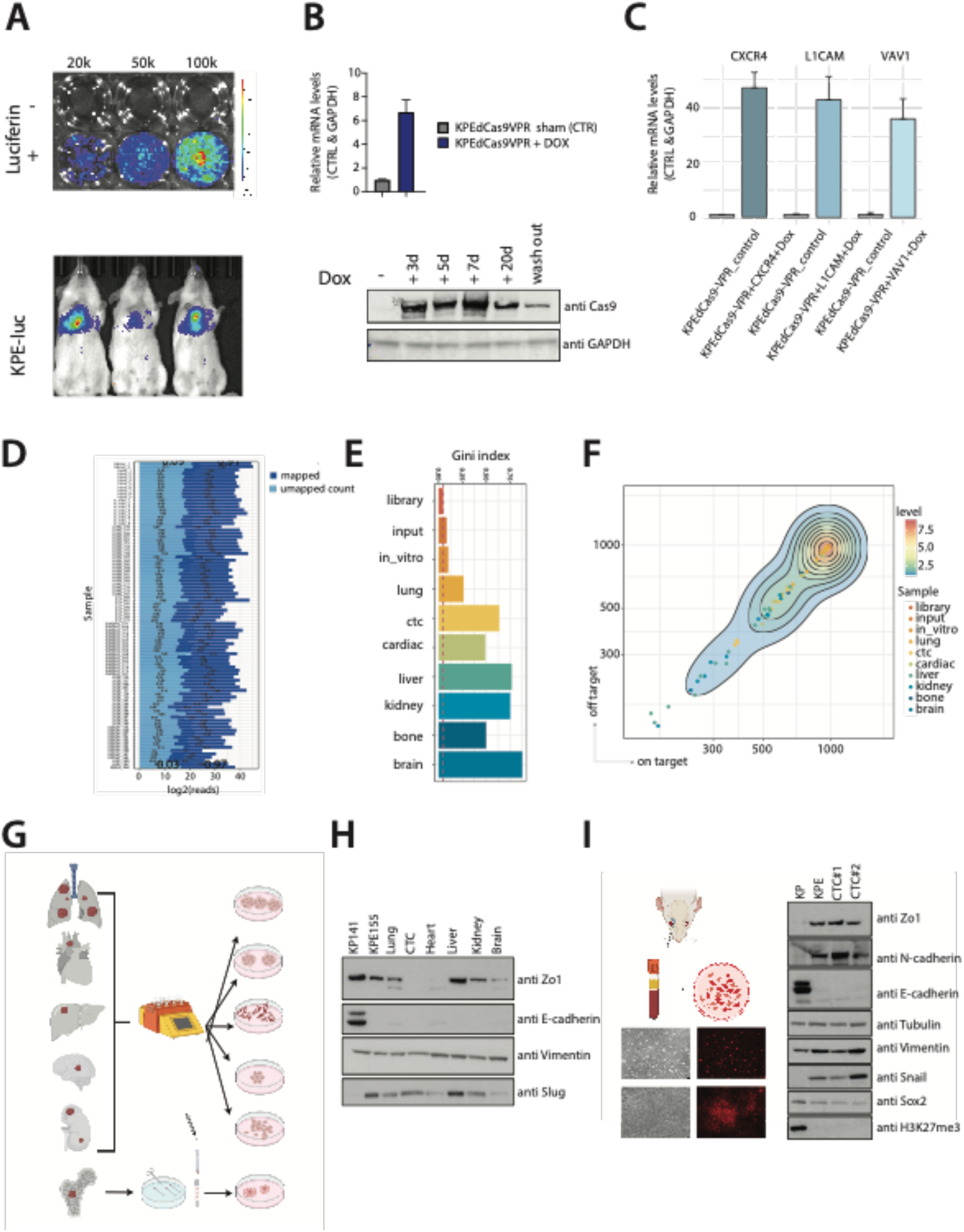
Extended characterization of the KPE-luc-tet-ON-dcas9VPR cellular system and in vivo CROP-seq technology. **A)** Above, Luciferase emission of *in vitro* KPE-cells treated with or without luciferin previously transduced with a vector containing Luciferase. Below, BLI upon grafting. Note the dose-response to luciferin. **B)** Bar plot showing *In vitro* expression by RT-qPCR (above) and western blot (below) of dcas9-vpr upon doxycycline treatment of KPE transduced with TetON-dcas9vpr system. **C)** Bar plot showing target gene expression detected by qPCR upon doxycycline treatment of *L1cam*, *Cxcr4*, and *Vav1* sgRNAs cloned in a CROP-seq vector. **D)** Bar plot showing reads mapping statistics **E)** Bar plot showing the distribution of the sgRNA evenness/GINI index across the indicated sample groups. **F)** Density plot showing the representation of on-target sgRNAs and off-target for the indicated tissues (color-coded). Note the overall linear correlation. **G)** Graphical depiction of the distinct isolation of primary tumor cells from tissues using gentleMACS™ Octo Dissociator with Heaters (Milteny) and from bone marrow cells using standard flushing. **H-I**) Western blot analysis of the indicated markers for the epithelial and mesenchymal phenotype in primary cells isolated from different tissues of mouse #60.

**Figure S2.**
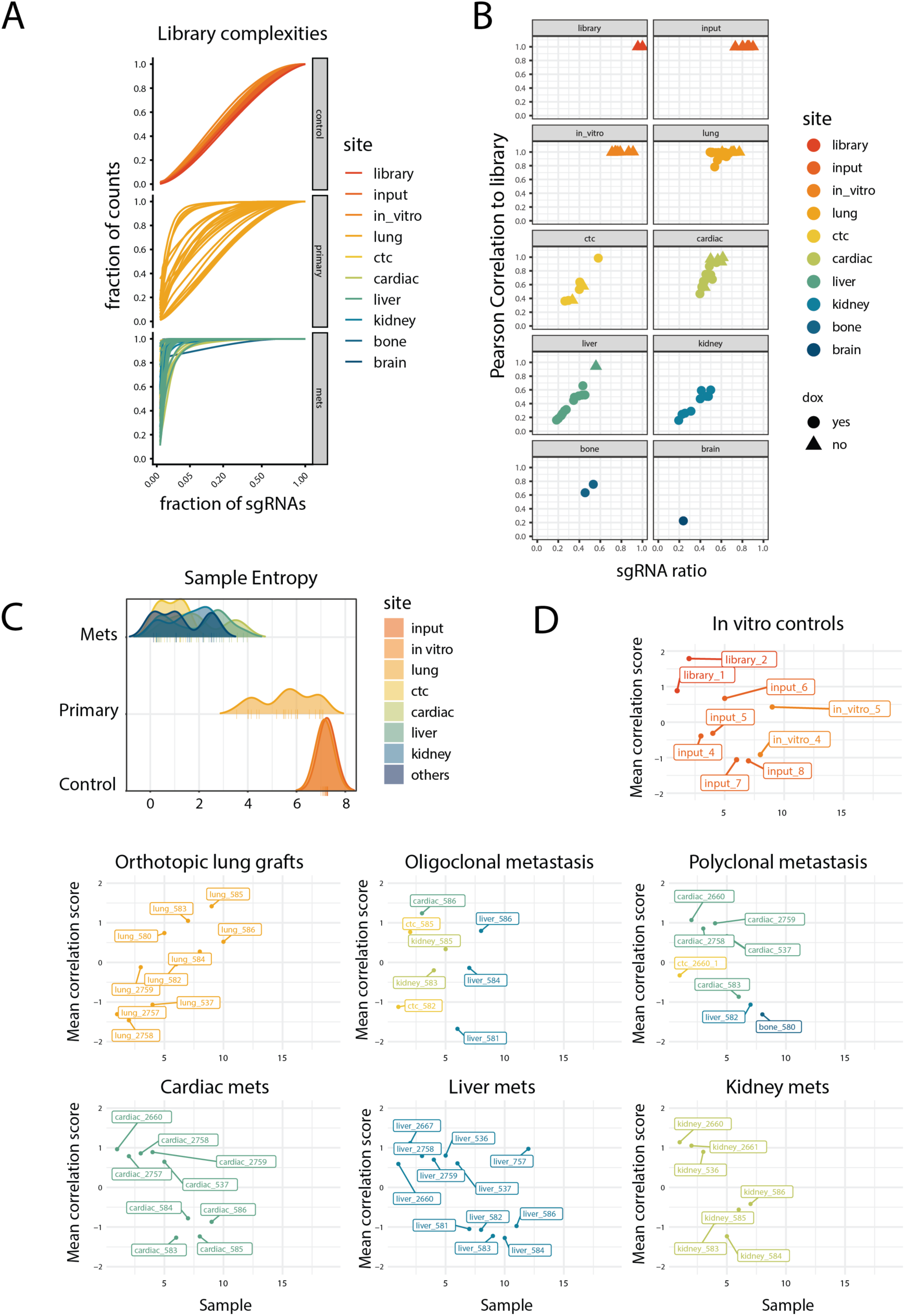
Extended characterization of the in vivo CRISPRa screen outcome. **A)** Library complexity distribution plot of the indicated sample groups. **B)** Scatter plot of the indicated sample scores subdivided by tissue of origin. **C)** Density distribution plot of sample entropy for the indicated groups. **D)** Scatter plot showing the distribution of the inter-group correlation scores for the indicated groups.

**Figure S3.**
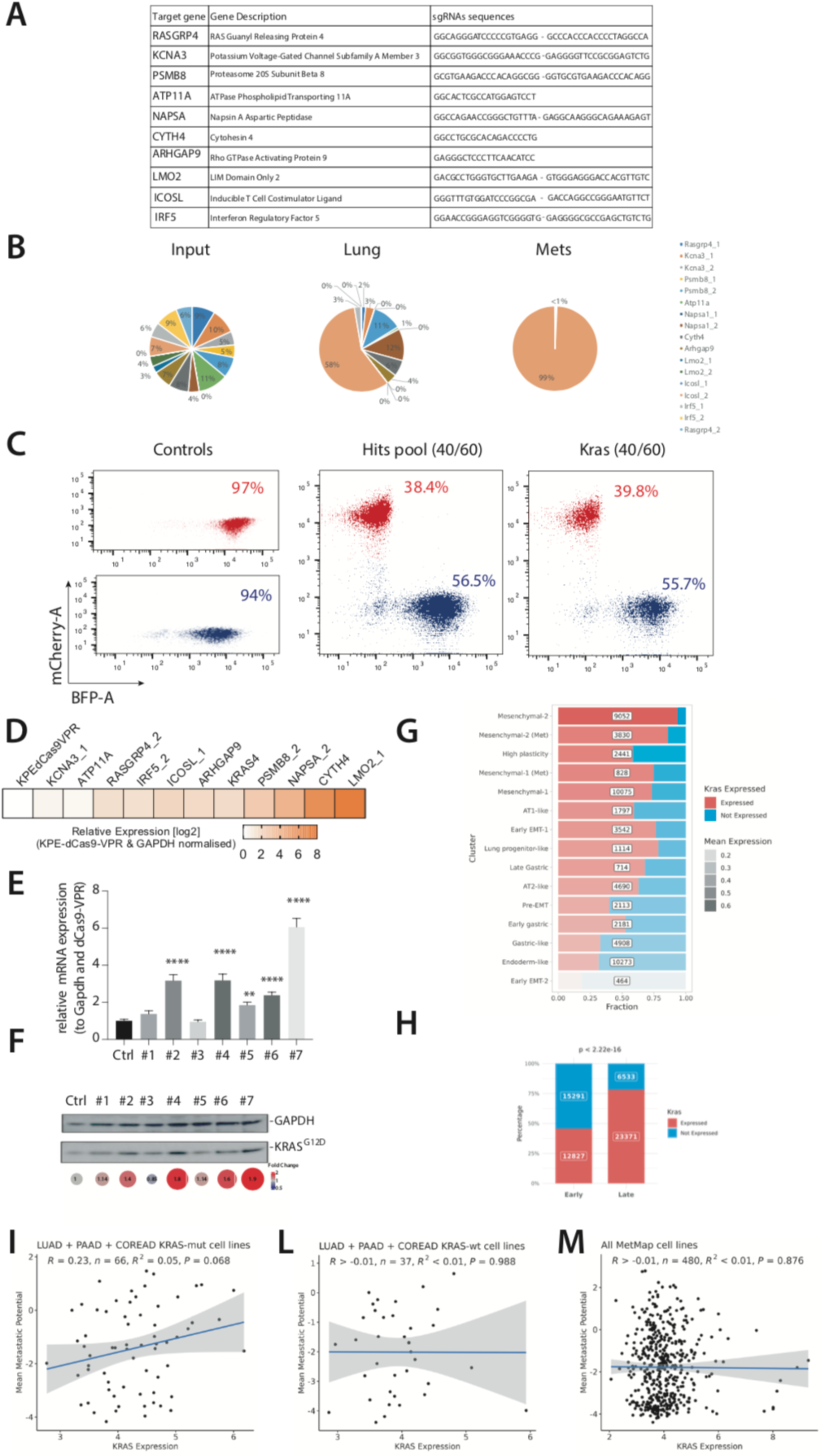
On-set *in vitro* specificity of KRAS and single genes overexpression and *in vivo* validation of the screening **A)** Table indicating the single gene description and the sgRNA sequences cloned in the CROP-seq-cherry vector used in the validation experiments **B)** pie chart indicating the frequency (%) of sgRNAs retrieved in the input, primary lung tumors, and metastatic organs in the in vivo validation screening of eight genes. **C)** FACS analysis of the in vivo competition experiment showing the injected population of BFP-nontargeting KPE input cells and Cherry-KPE targeting cells. **D)** Heat map depiction of quantitative PCR (qPCR) data. Each single gene expression in KPE was assessed upon dox-inducible dcas9-vpr activation after 7 or 14 days. Data are normalized by gapdh and dCas9-VPR, and p-values are by 1-way ANOVA **E)** Expression level measurement upon doxycycline treatment of KPE cells transduced with seven single sgRNAs targeting KRAS. Bar plot showing mRNA levels referring to GAPDPH and control KPE. **F)** western blot, and bubble blot showing KRASG12D protein levels and the respective quantification (right). **G)** Horizontal stacked bar plot showing the number of single cells from autochthonous KP- tracer mice in which *Kras* expression is detected or not, as proxy for high and low expression, respectively. Cells are grouped according to their cell state as originally defined by Yang et al., 2022. **H)** Vertical stacked bar plot showing the number of single cells from autochthonous KP- tracer mice in which *Kras* expression is detected or not, as proxy for high and low expression, respectively. Cells are grouped according to their progression stage as originally defined by Yang et al., 2022 and significance is calculated by Fisher’s exact test. **I-M**) Scatter plot showing the correlation between KRAS expression and metastatic potential in recipient mice as defined by MetMap500 in Jin et al., 2020, for the indicated subsets of barcoded CCLE cell lines. Pearson’s R and P value of correlation are shown. Note the positive correlation for KRAS expression limited to cell lines derived from KRAS-driven cancers with KRAS driver mutations.

**Figure S4.**
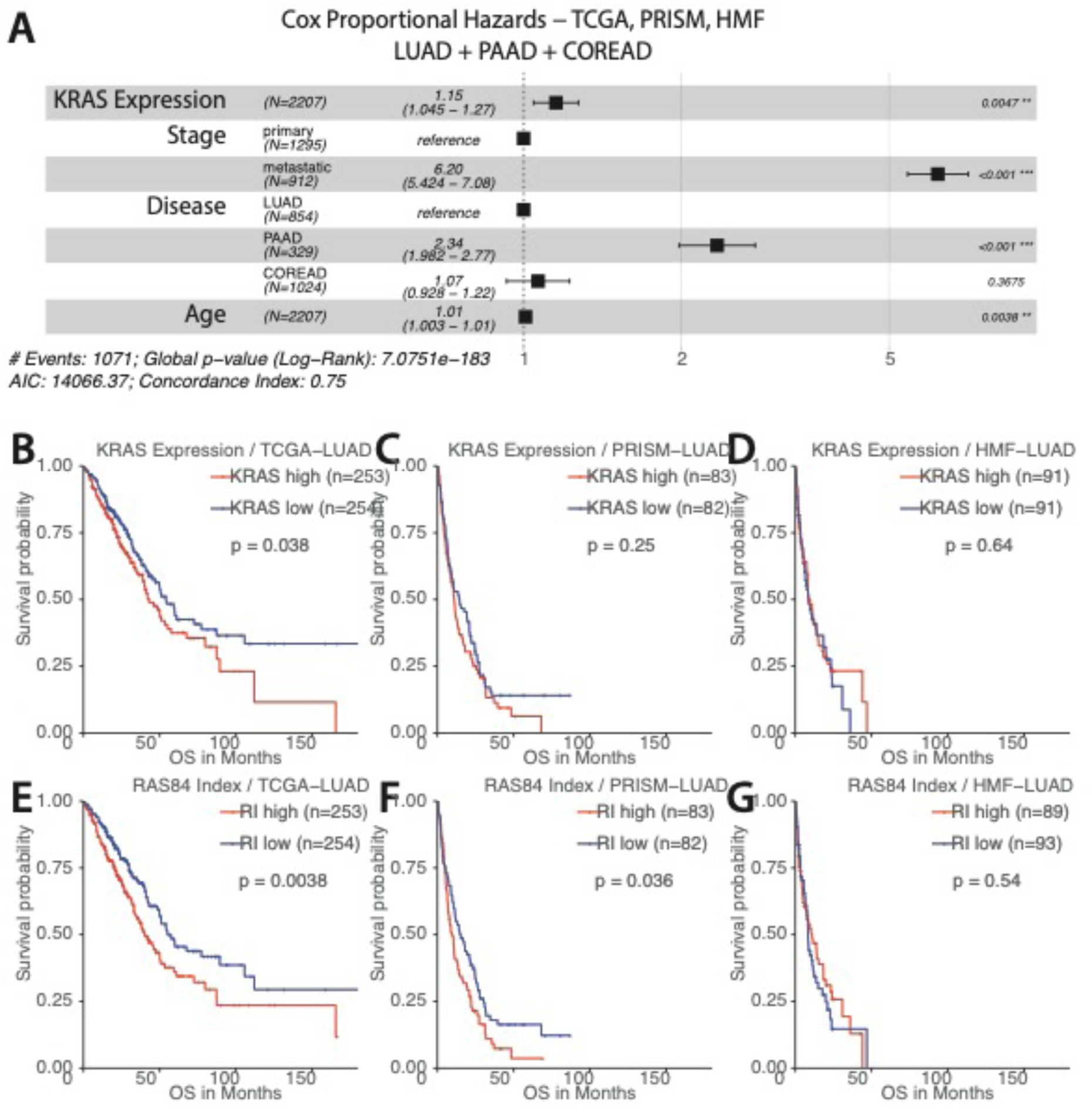
KRAS transcriptional levels are linked to poor survival in Kras-driven cancer **A)** Forest plot of the Cox proportional hazards model illustrating the effect of KRAS expression (see methods) on survival in LUAD, PAAD, COREAD patient cohorts from TCGA, PRISM and HMF. KRAS expression is adjusted for stage and age. P-value and hazard ratio of a covariate is indicated in the right and middle of a row respectively. Asterisks denotes significance. Global p-value is indicated in the bottom. **B-E)** Kaplan–Meier survival plots and Log-rank test for overall survival in primary (TCGA) and metastatic (PRISM and HMF) cohorts. Samples were separated into groups based on median KRAS gene expression level (see methods). **F-I**) Kaplan–Meier survival plots and Log-rank test for the same comparisons but focused on the transcriptional signature optimised to capture RAS oncogenic activity (RAS84 Index, RI) in LUAD (East et al). Samples were separated into groups based on median RI level (see methods).

**Figure S5.**
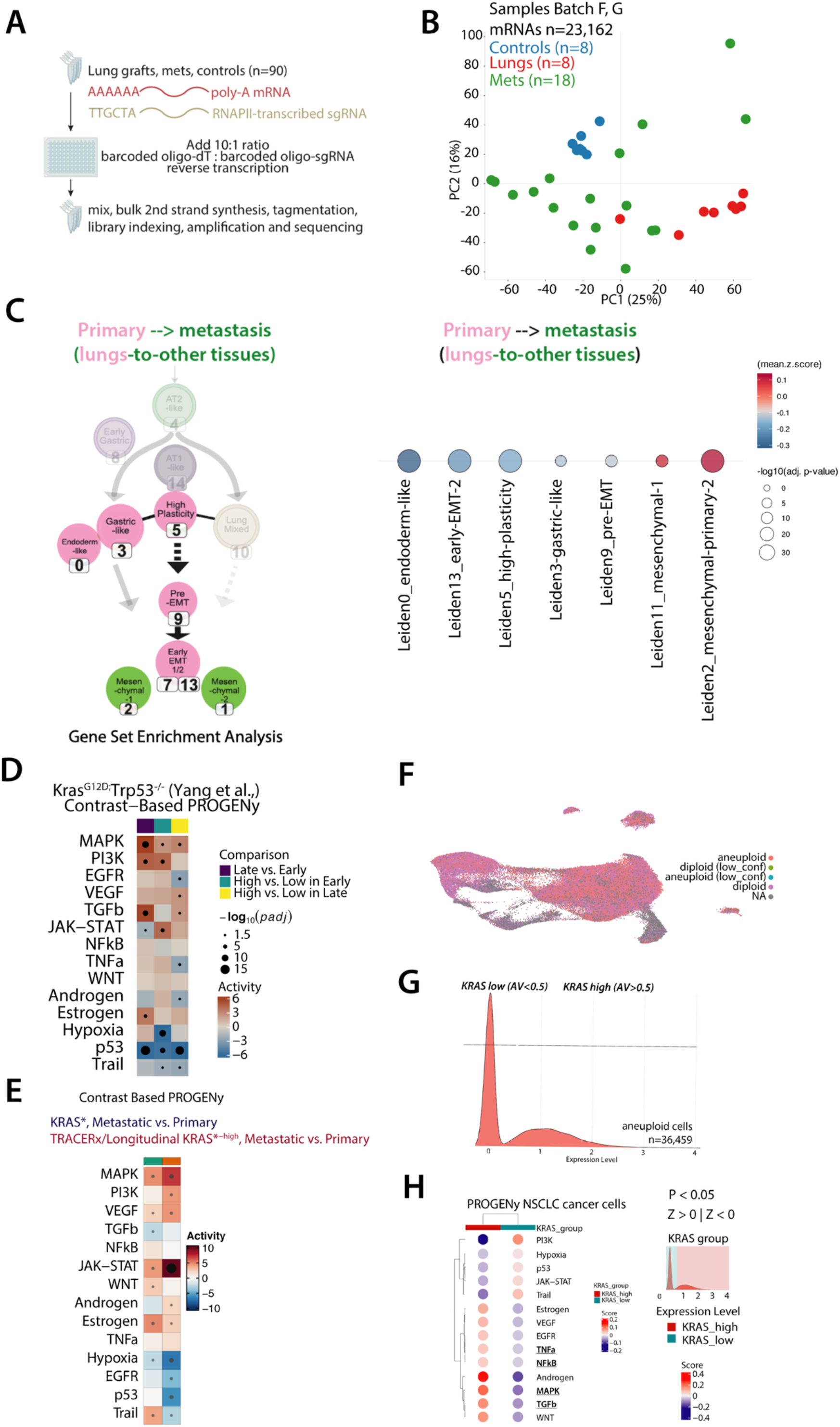
Extended analysis on transcriptomic analyses. **A)** Workflow integrating CROP-seq and BRB-seq to capture targeting gRNAs, non-targeting cellular barcodes and global poly-A mRNA. **B)** Representative hierarchical clustering of Euclidean distances between the indicated RNA-seq samples. **C)** Right, graphical representation of GSEA of lung differential expression between lung and metastatic samples from mRNA-seq. Color code and position in the evolutionary plot and dot size denote the directionality of the GSEA and gene set size, respectively. Left, schematic depiction of GSEA in the cell states associated with autochthonous KP-tracer model LUAD progression as defined by scRNA-seq and lineage-tracing (Yang et al., 2022). **D)** Heatmap showing activity scores (presented as z-score) of PROGENy pathways in single cell clusters associated with progression stages and separated based on Kras expression as in Fig. 2F from the autochthonous KP-tracer model as defined by scRNA-seq and lineage-tracing (Yang et al., 2022). **E)** Heatmap showing activity scores (presented as z-score) of PROGENy pathways in n=1467 patients with LUAD from primary (TCGA), and metastatic cohorts (PRISM and Hartwig) as well as longitudinal TRACERx. **F)** UMAP of scRNA-seq from 83,701 NSCLC cells across 42 patients samples and classified by aneuploidy estimation using CopyKAT. **G)** Density plot of normalized *KRAS* expression level in 36,459 cells aneuplouid cells from (F). **H)** Dotplot plot heatmap showing activity scores (presented as z-score) of PROGENy pathways (left) in NSCLC patients (Wu et al., n=42 patients, aneuploid cells n=83,701 cells)

**Figure S6.**
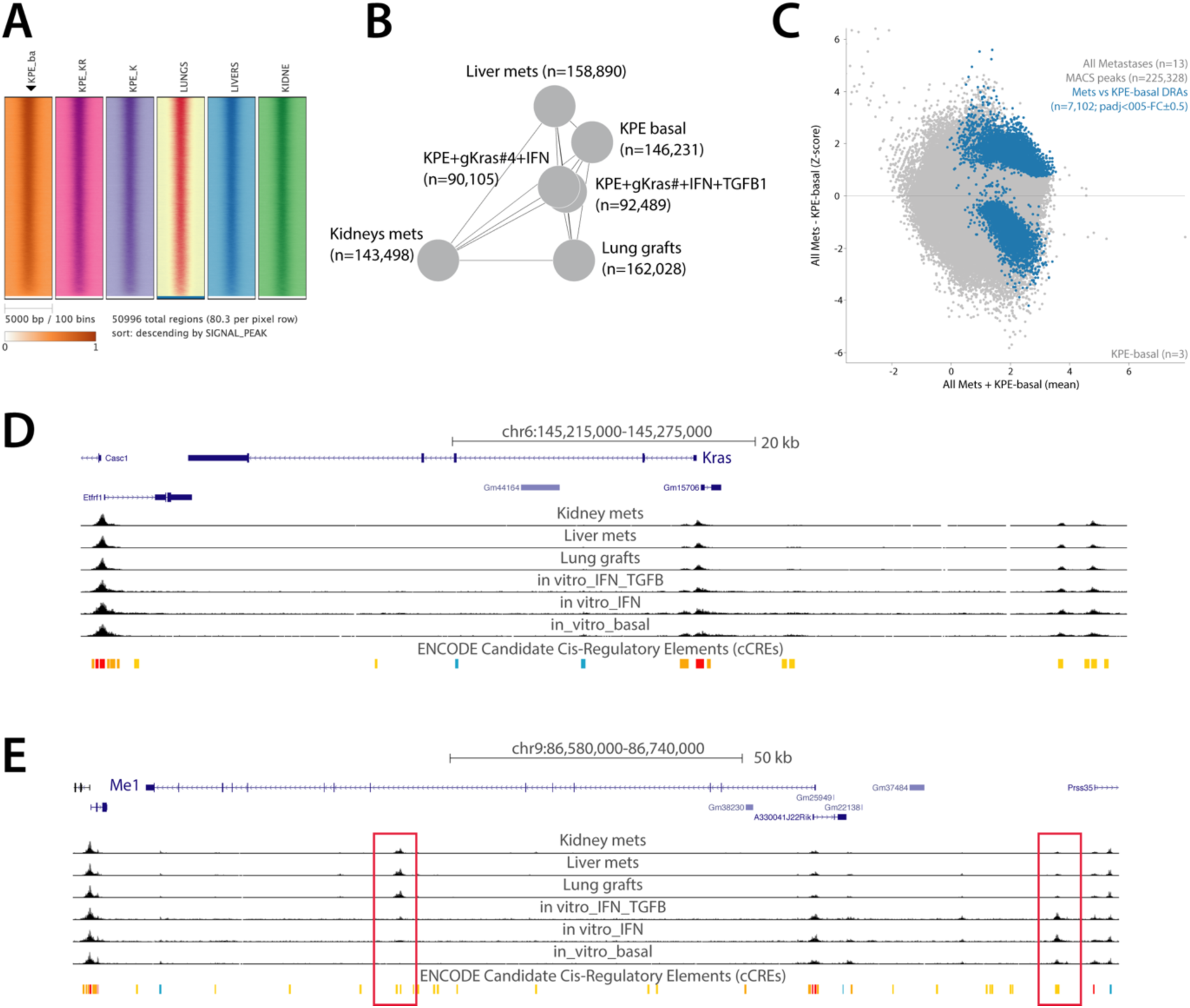
Extended analysis of integration of Kras with chromatin in KPE and its significance in LUAD patients’ datasets. **A)** Heatmap of the indicated ATAC-seq (merged) tracks for the shared peaks from the upset plot in 6b. **B)** Giraph plot showing the distance between individual ATAC-seq peak lists. Note the quantitative difference for the in vivo and in vitro chromatin accessibility despite the qualitative overlap in 6b and s6a. **C)** MA plot showing distribution of the indicated ATAC-seq DAR (blue) compared to all other accessible peaks (grey). **D-E**) UCSC view of Kras and Me1 to illustrate the quality of the ATAC-seq profile, the shared peaks and the ones private to progression such as the gene set in s6c. The red boxes denote opposite behaviour in accessibility.

**Figure S7.**
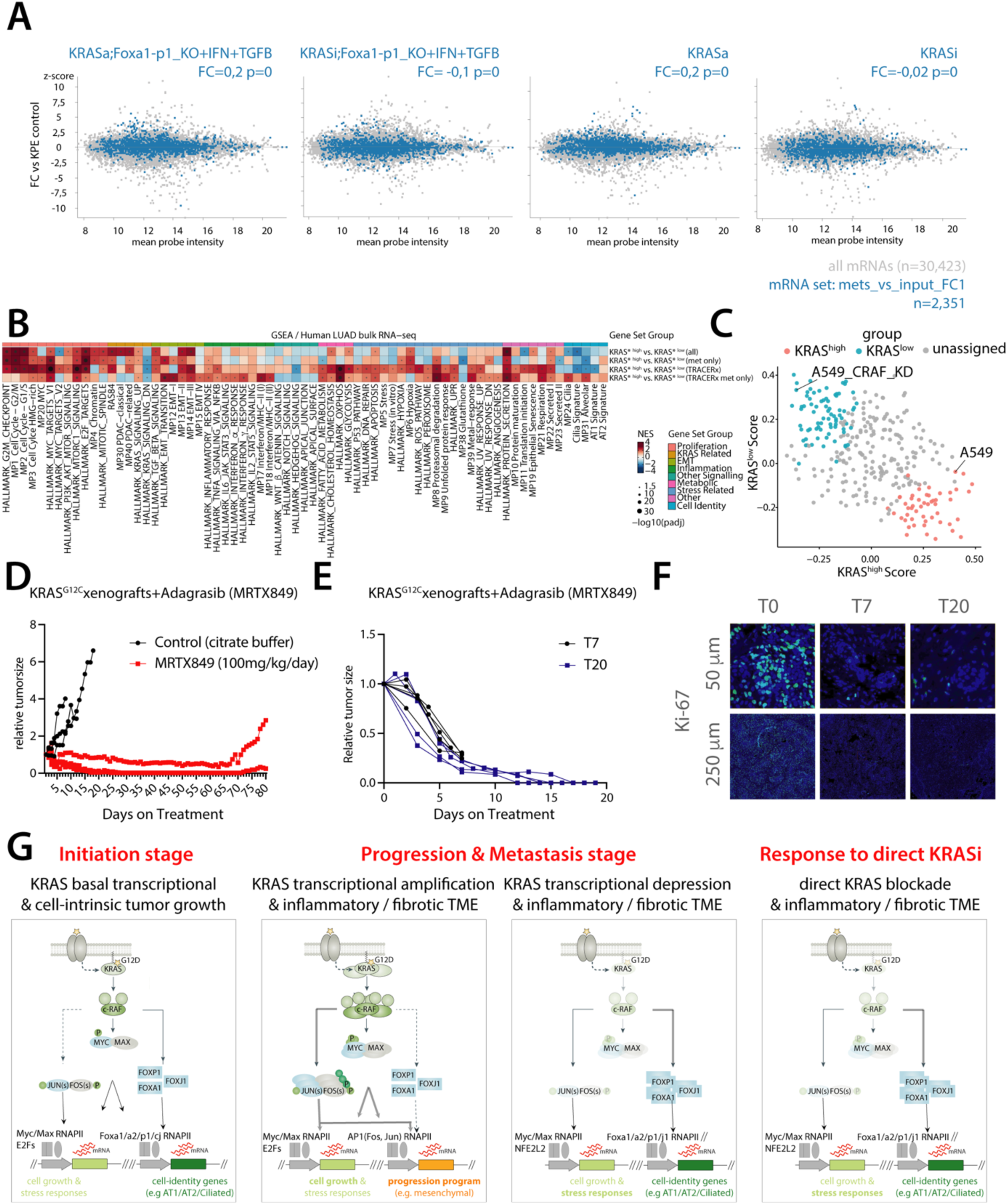
Extended transcriptomic analyses and KRAS dosage model. **A)** MA plot and GSEA showing distribution of the gene residing in chromatin remodelled in KPE in vivo (blue; **Fig. 6B-6D**) and the directionality, extent and significance of their enrichment as compared to all other mRNAs (grey). **B)** Extended GSEA scores (presented as normalized enrichment scores) of the indicated pan-cancer single-cell consensus meta-programs and selected hallmarks in KRAS high vs. low contrast setting (see methods) in primary & metastatic bulk human RNA-seq profiles. Patients are divided according to their known KRAS mutation status (“*” legend). Filled circles denote Benjamini-Hochberg adjusted p-value level: *, p < 0.05; **, p < 0.01; ***, p < 0.001. **C)** Scatter plot showing the distribution of CCLE cell lines from the DepMap according to their enrichment score for the human KRAS high and low signature, X and Y axiss, respectively. Color code denotes their classification as KRAS high (pink), low (blue) or unclassified (grey) based on thresholds defined as described in the methods. **D-E)** Line plots showing the longitudinal tumor size measuring over time for human KRAS^G12C^ PDX models. MRTX849 was administered via daily oral via IP at the indicated dose to mice bearing LUAD xenografts stabilized in four previous passages (F4). Dosing was initiated when tumors were approximately 300 mm^3^. MRTX849 was administered to mice daily until day 67. Individual data point denote a single tumor volume per mouse. Long-term and short term sampling is shown in D and E, respectively. **F)** Representative immunohistochemical (IHC) images of PDX tumors collected at indicated timepoints, stained for Ki-67 (green), and counterstained with DAPI (blue). Scale size is indicated. **G)** A model for *KRAS*-dosage driven tumor evolution and LUAD cell states. *KRAS* gene level are normal at the initiation stages (e.g. AT2 transformed cells) and KRAS signal transduction promotes the transcription of both cell proliferation and cell identity genes. During progression & metastasis, LUAD cells experiencing pressure to increase *KRAS* transcriptional dosage promotes AP-1 accumulation and activity at genes involved in progression and metastasis (e.g. EMT genes driven by pro-inflammatory/pro-fibrotic TME).

## References

1. Pérez-González, A., Bévant, K. & Blanpain, C. Cancer cell plasticity during tumor progression, metastasis and response to therapy. Nat Cancer 4, 1063–1082 (2023).

2. Altorki, N. K. et al. The lung microenvironment: an important regulator of tumour growth and metastasis. Nat Rev Cancer 1–23 (2018). doi:10.1038/s41568-018-0081-9

3. Nguyen, D. X., Bos, P. D. & Massagué, J. Metastasis: from dissemination to organ-specific colonization. Nat Rev Cancer 9, 274–284 (2009).

4. Pradat, Y. et al. Integrative pan-cancer genomic and transcriptomic analyses of refractory metastatic cancer. Cancer Discovery (2023). doi:10.1158/2159-8290.CD-22-0966

5. Priestley, P. et al. Pan-cancer whole-genome analyses of metastatic solid tumours. Nature 575, 210–216 (2019).

6. Karras, P., Black, J. R. M., McGranahan, N. & Marine, J.-C. Decoding the interplay between genetic and non-genetic drivers of metastasis. Nature 629, 543–554 (2024).

7. Gargiulo, G., Serresi, M. & Marine, J.-C. Cell States in Cancer: Drivers, Passengers, and Trailers. Cancer Discovery 14, 610–614 (2024).

8. Ostrem, J. M., Peters, U., Sos, M. L., Wells, J. A. & Shokat, K. M. K-Ras(G12C) inhibitors allosterically control GTP affinity and effector interactions. Nature 503, 548–551 (2013).

9. Wasko, U. N. et al. Tumor-selective activity of RAS-GTP inhibition in pancreatic cancer. Nature 1–3 (2024). doi:10.1038/s41586-024-07379-z

10. Holderfield, M. et al. Concurrent inhibition of oncogenic and wild-type RAS-GTP for cancer therapy. Nature 1–8 (2024). doi:10.1038/s41586-024-07205-6

11. Jiang, J. et al. Translational and Therapeutic Evaluation of RAS-GTP Inhibition by RMC-6236 in RAS-Driven Cancers. Cancer Discovery OF1–OF24 (2024). doi:10.1158/2159-8290.CD-24-0027

12. Cancer Genome Atlas Research Network. Comprehensive molecular profiling of lung adenocarcinoma. Nature 511, 543–550 (2014).

13. East, P. et al. RAS oncogenic activity predicts response to chemotherapy and outcome in lung adenocarcinoma. Nat Commun 13, 1–17 (2022).

14. Vogelstein, B. et al. Genetic Alterations during Colorectal-Tumor Development. N Engl J Med 319, 525–532 (1988).

15. Jackson, E. L. et al. Analysis of lung tumor initiation and progression using conditional expression of oncogenic K-ras. Genes Dev 15, 3243–3248 (2001).

16. Tuveson, D. A. et al. Endogenous oncogenic K-ras(G12D) stimulates proliferation and widespread neoplastic and developmental defects. Cancer Cell 5, 375–387 (2004).

17. Li, Z. et al. Alveolar Differentiation Drives Resistance to KRAS Inhibition in Lung Adenocarcinoma. Cancer Discovery 14, 308–325 (2024).

18. Yang, D. et al. Lineage tracing reveals the phylodynamics, plasticity, and paths of tumor evolution. Cell 185, 1905–1923.e25 (2022).

19. Lafave, L. M. et al. Epigenomic State Transitions Characterize Tumor Progression in Mouse Lung Adenocarcinoma. Cancer Cell (2020). doi:10.1016/j.ccell.2020.06.006

20. Serresi, M. et al. Polycomb Repressive Complex 2 Is a Barrier to KRAS-Driven Inflammation and Epithelial-Mesenchymal Transition in Non-Small-Cell Lung Cancer. Cancer Cell 29, 17–31 (2016).

21. Caswell, D. R. et al. Obligate progression precedes lung adenocarcinoma dissemination. Cancer Discovery 4, 781–789 (2014).

22. Winslow, M. M. et al. Suppression of lung adenocarcinoma progression by Nkx2-1. Nature 473, 101–104 (2011).

23. Sutherland, K. D. et al. Multiple cells-of-origin of mutant K-Ras-induced mouse lung adenocarcinoma. Proc Natl Acad Sci USA (2014). doi:10.1073/pnas.1319963111

24. Kerk, S. A., Papagiannakopoulos, T., Shah, Y. M. & Lyssiotis, C. A. Metabolic networks in mutant KRAS-driven tumours: tissue specificities and the microenvironment. Nat Rev Cancer 21, 510–525 (2021).

25. Najumudeen, A. K. et al. The amino acid transporter SLC7A5 is required for efficient growth of KRAS-mutant colorectal cancer. Nat Genet 53, 16–26 (2021).

26. Mueller, S. et al. Evolutionary routes and KRAS dosage define pancreatic cancer phenotypes. Nature Publishing Group 1–28 (2018). doi:10.1038/nature25459

27. Kerr, E. M., Gaude, E., Turrell, F. K., Frezza, C. & Martins, C. P. Mutant Kras copy number defines metabolic reprogramming and therapeutic susceptibilities. Nature 531, 110– 113 (2016).

28. Serresi, M. et al. Ezh2 inhibition in Kras-driven lung cancer amplifies inflammation and associated vulnerabilities. J Exp Med 215, 3115–3135 (2018).

29. Gargiulo, G., Serresi, M., Cesaroni, M., Hulsman, D. & Van Lohuizen, M. In vivo shRNA screens in solid tumors. Nat Protoc 9, 2880–2902 (2014).

30. Camolotto, S. A., et al. FoxA1 and FoxA2 drive gastric differentiation and suppress squamous identity in NKX2-1-negative lung cancer. Elife 7, (2018).

31. Araujo, H. A. et al. Mechanisms of response and tolerance to active RAS inhibition in KRAS-mutant NSCLC. Cancer Discovery (2024). doi:10.1158/2159-8290.CD-24-0421

32. Chavez, A. et al. Highly efficient Cas9-mediated transcriptional programming. Nat Methods 12, 326–328 (2015).

33. Datlinger, P. et al. Pooled CRISPR screening with single-cell transcriptome readout. Nat Methods 14, 297–301 (2017).

34. Frankell, A. M. et al. The evolution of lung cancer and impact of subclonal selection in TRACERx. Nature 616, 525–533 (2023).

35. Laughney, A. M. et al. Regenerative lineages and immune-mediated pruning in lung cancer metastasis. Nat Med 26, 259–269 (2020).

36. Wu, F. et al. Single-cell profiling of tumor heterogeneity and the microenvironment in advanced non-small cell lung cancer. Nat Commun 12, 2540–11 (2021).

37. Downward, J. Targeting RAS signalling pathways in cancer therapy. Nat Rev Cancer 3, 11–22 (2003).

38. Eferl, R. & Wagner, E. F. AP-1: a double-edged sword in tumorigenesis. Nat Rev Cancer 3, 859–868 (2003).

39. Sekiya, T., Muthurajan, U. M., Luger, K., Tulin, A. V. & Zaret, K. S. Nucleosome-binding affinity as a primary determinant of the nuclear mobility of the pioneer transcription factor FoxA. Genes Dev 23, 804–809 (2009).

40. Gavish, A. et al. Hallmarks of transcriptional intratumour heterogeneity across a thousand tumours. Nature 618, 598–606 (2023).

41. Hallin, J. et al. The KRASG12C Inhibitor MRTX849 Provides Insight toward Therapeutic Susceptibility of KRAS-Mutant Cancers in Mouse Models and Patients. Cancer Discovery 10, 54–71 (2020).

42. Hahn, W. C. et al. Creation of human tumour cells with defined genetic elements. Nature 400, 464–468 (1999).

43. Zafra, M. P. et al. An In Vivo Kras Allelic Series Reveals Distinct Phenotypes of Common Oncogenic Variants. Cancer Discovery 10, 1654–1671 (2020).

44. To, M. D. et al. Kras regulatory elements and exon 4A determine mutation specificity in lung cancer. 40, 1240–1244 (2008).

45. Ambrogio, C. et al. KRAS Dimerization Impacts MEK Inhibitor Sensitivity and Oncogenic Activity of Mutant KRAS. Cell 172, 857–868.e15 (2018).

46. Yan, H. et al. Loss of the wild-type KRAS allele promotes pancreatic cancer progression through functional activation of YAP1. Oncogene 40, 6759–6771 (2021).

47. Najumudeen, A. K. et al. KRAS allelic imbalance drives tumour initiation yet suppresses metastasis in colorectal cancer in vivo. Nat Commun 15, 100–14 (2024).

48. Bakir, Al, M., et al. The evolution of non-small cell lung cancer metastases in TRACERx. Nature 616, 534–542 (2023).

49. Ivanisevic, T. et al. Increased dosage of wild-type KRAS protein drives KRAS-mutant lung tumorigenesis and drug resistance. bioRxiv 2024.02.27.582346 (2024). doi:10.1101/2024.02.27.582346

50. Tape, C. J. et al. Oncogenic KRAS Regulates Tumor Cell Signaling via Stromal Reciprocation. Cell 165, 910–920 (2016).

51. Jenks, A. D. et al. Primary Cilia Mediate Diverse Kinase Inhibitor Resistance Mechanisms in Cancer. Cell Reports 23, 3042–3055 (2018).

52. Chen, F. et al. Polycomb deficiency drives a FOXP2-high aggressive state targetable by epigenetic inhibitors. Nat Commun 14, 336–18 (2023).

53. Kim, M. P. et al. Oncogenic KRAS Recruits an Expansive Transcriptional Network through Mutant p53 to Drive Pancreatic Cancer Metastasis. Cancer Discovery 11, 2094–2111 (2021).

54. Li, Y. et al. Mutant Kras co-opts a proto-oncogenic enhancer network in inflammation-induced metaplastic progenitor cells to initiate pancreatic cancer. Nat Cancer 1–45 (2021). doi:10.1038/s43018-020-00134-z

55. Ying, H. et al. Oncogenic Kras Maintains Pancreatic Tumors through Regulation of Anabolic Glucose Metabolism. Cell 149, 656–670 (2012).

56. Viale, A. et al. Oncogene ablation-resistant pancreatic cancer cells depend on mitochondrial function. Nature (2014). doi:10.1038/nature13611

57. Biddie, S. C. et al. Transcription factor AP1 potentiates chromatin accessibility and glucocorticoid receptor binding. Mol Cell 43, 145–155 (2011).

58. Gargiulo, G. et al. In vivo RNAi screen for BMI1 targets identifies TGF-β/BMP-ER stress pathways as key regulators of neural-and malignant glioma-stem cell homeostasis. Cancer Cell 23, 660–676 (2013).

59. Zhao, Y. et al. ‘Stripe’ transcription factors provide accessibility to co-binding partners in mammalian genomes. Mol Cell 82, 3398–3411.e11 (2022).

60. Klomp, J. A. et al. Defining the KRAS- and ERK-dependent transcriptome in KRAS-mutant cancers. Science 384, eadk0775 (2024).

61. Klomp, J. E. et al. Determining the ERK-regulated phosphoproteome driving KRAS-mutant cancer. Science 384, eadk0850 (2024).

62. Datlinger, P. et al. Ultra-high-throughput single-cell RNA sequencing and perturbation screening with combinatorial fluidic indexing. Nat Methods 18, 635– 642 (2021).

63. Alpern, D. et al. BRB-seq: ultra-affordable high-throughput transcriptomics enabled by bulk RNA barcoding and sequencing. 1–15 (2019). doi:10.1186/s13059-019-1671-x

## Methods References

Love MI, Huber W, Anders S. Moderated estimation of fold change and dispersion for RNA-seq data with DESeq2. _Genome Biol_. 2014;15(12):550. doi:[10.1186/s13059-014-0550-8](10.1186/s13059-014-0550-8)

Hänzelmann S, Castelo R, Guinney J. GSVA: gene set variation analysis for microarray and RNA-seq data. _BMC Bioinformatics_. 2013;14:7. doi:[10.1186/1471-2105-14- 7](10.1186/1471-2105-14-7)

Bhuva DD, Cursons J, Davis MJ. Stable gene expression for normalisation and single-sample scoring. _Nucleic Acids Res_. 2020;48(19):e113. doi:[10.1093/nar/gkaa802](10.1093/nar/gkaa802)

Zhang Y, Parmigiani G, Johnson WE. ComBat-seq: batch effect adjustment for RNA-seq count data. _NAR Genom Bioinform_. 2020;2(3):lqaa078. doi:[10.1093/nargab/lqaa078](10.1093/nargab/lqaa078)

East P, Kelly GP, Biswas D, et al. RAS oncogenic activity predicts response to chemotherapy and outcome in lung adenocarcinoma. _Nat Commun_. 2022;13(1):5632. doi:[10.1038/s41467-022-33290-0](10.1038/s41467-022-33290-0)

Korotkevich G, Sukhov V, Budin N, Shpak B, Artyomov MN, Sergushichev A. Fast gene set enrichment analysis. Published online February 1, 2021:060012. doi:[10.1101/060012](10.1101/060012)

Badia-I-Mompel P, Vélez Santiago J, Braunger J, et al. decoupleR: ensemble of computational methods to infer biological activities from omics data. _Bioinform Adv_. 2022;2(1):vbac016. doi:[10.1093/bioadv/vbac016](10.1093/bioadv/vbac016)

Schubert M, Klinger B, Klünemann M, et al. Perturbation-response genes reveal signaling footprints in cancer gene expression. _Nat Commun_. 2018;9(1):20. doi:[10.1038/s41467-017-02391-6](10.1038/s41467-017-02391-6)

Müller-Dott S, Tsirvouli E, Vazquez M, et al. Expanding the coverage of regulons from high-confidence prior knowledge for accurate estimation of transcription factor activities. _Nucleic Acids Res_. Published online October 16, 2023:gkad841. doi:[10.1093/nar/gkad841](10.1093/nar/gkad841)

Tsherniak A, Vazquez F, Montgomery PG, et al. Defining a Cancer Dependency Map. _Cell_. 2017;170(3):564–576.e16. doi:[10.1016/j.cell.2017.06.010](10.1016/j.cell.2017.06.010)

Jin X, Demere Z, Nair K, et al. A metastasis map of human cancer cell lines. _Nature_. 2020;588(7837):331-336. doi:[10.1038/s41586-020-2969-2](10.1038/s41586-020-2969-2)

